# Chemical Descriptors and Deep Learning Embeddings for Scoring *de novo* Peptide Designs

**DOI:** 10.64898/2026.09.10.750670

**Authors:** Quentin Trolliet, Alex Abrudan, Aryan Bhasin, Yunguan Fu, Joao Paulo Euko, Cheng Zhang, Francesco Saccon

**Affiliations:** InstaDeep Ltd, 1 Triton Square, London NW1 3BF, London, United Kingdom; Yusuf Hamied Department of Chemistry, University of Cambridge, Cambridge, UK; Nxera Pharma; Boltz PBC

**Keywords:** Protein stability, Deep learning, Protein language models, Peptide design, Binding affinity

## Abstract

Peptides occupy a valuable niche between small molecules and biologics, but the clinical translation of *de novo* peptide designs requires rigorous scoring to simultaneously optimise target binding affinity alongside multiple developability traits, including stability, membrane permeability, aggregation propensity, and non-fouling behaviour. Here, we evaluate two distinct approaches for scoring these candidates: classical chemical descriptors and modern deep learning representations derived from protein language and folding models. Assembling nine public datasets spanning five developability traits and four binding-affinity endpoints, we find sequence-derived chemical descriptors alone contain sufficient information to predict developability task labels effectively. Given their drastically lower computational cost and higher interpretability, classical machine learning models trained on these simple descriptors frequently match or approach the performance of complex deep learning architectures, emerging as a highly efficient and interpretable alternative for high-throughput scoring. Finally, for scoring binding affinity, we demonstrate that Boltz-2 pair representations capture the most information among the tested representations; however, the model’s predictive power is confounded by a significant bias from the molecular weight of the peptides. Together, these results establish a comprehensive assessment of state-of-the-art methods for predicting both peptide developability and binding affinity, highlighting the enduring value of interpretable chemical descriptors alongside deep learning in the scoring and selection of *de novo* peptide designs.

## 1 Introduction

Peptides are rapidly emerging as highly valuable therapeutics for treating an array of metabolic diseases, oncological indications, and infectious disorders due to their remarkable target specificity, low tissue accumulation, and favourable safety profiles compared to synthetic small molecules [Muttenthaler et al., 2021]. Deep learning (DL) generative pipelines have opened unprecedented avenues for the *de novo* design of such therapeutics. Advanced frameworks, including Bindcraft [Pacesa et al., 2025], BoltzGen [Stark et al., 2025], RFpeptides [Rettie et al., 2025], and LatentX [Latent Labs Team et al., 2025], now allow researchers to rapidly sample vast regions of the chemical and structural space, generating massive libraries of novel peptide candidates tailored to specific biological targets. While these emerging generative pipelines excel at producing novel sequences, translating these computationally designed candidates into successful clinical therapeutics requires navigating a complex, hyper-dimensional developability landscape. Engineering a viable therapeutic peptide demands the simultaneous optimisation of target binding affinity, cellular membrane permeability for intracellular targets, proteolytic stability against rapid serum degradation, aqueous solubility, and scalable manufacturability [Rossino et al., 2023, Li et al., 2023]. Crucially, target binding affinity and developability attributes must be evaluated as deeply interconnected axes of the same optimisation problem [Hopkins et al., 2004]. An engineered peptide with exceptional, nanomolar target affinity remains clinically unviable if it concurrently exhibits severe systemic toxicity, poor proteolytic stability, or insurmountable manufacturing bottlenecks [Lau and Dunn, 2018].

DL models have become increasingly important to explore this vast combinatorial sequence space efficiently, shifting the paradigm from passive, resource-intensive experimental screening towards predictive and generative multi-property optimisation frameworks [Erckes et al., 2026]. Protein language models (PLM) are trained by masked language modelling over hundreds of millions of unaligned sequences, and their per-residue embeddings capture evolutionary and biophysical regularities without ever being shown a structure [Lin et al., 2023]. Co-folding models instead mine co-evolutionary signal from a multiple sequence alignment into a pairwise representation over token pairs, from which a separate module generates coordinates [Jumper et al., 2021, Passaro et al., 2025]. To harness these capabilities for therapeutic discovery, several recent architectures have adapted these generalized protein embeddings for peptide-specific endpoints. For instance, PeptideBERT fine-tunes the ProtBERT language model to predict hemolysis, solubility, and non-fouling [Guntuboina et al., 2023]. Other groups have instead represented peptides as molecular graphs and used them to train graph neural networks: PepLand pre-trains across atom and fragment views to handle both canonical and non-canonical amino acids [Zhang et al., 2025], and pepADMET combines molecular graphs with enzymatic descriptors across nineteen ADMET endpoints [Tan et al., 2026]. Furthermore, co-folding models have recently been extended with small molecule binding affinity prediction heads [Passaro et al., 2025, Shenoy et al., 2026, SandboxAQ, 2026], but this approach remains unevaluated for peptides.

Therefore, a fundamental question remains: which features are most informative for these predictive tasks, and which representations best capture them? Historically, physicochemical properties derived directly from the primary amino acid sequence have provided a valuable toolkit for identifying high-level functional and developability traits [Ofer and Linial, 2015], offering a high degree of biological interpretability. Sequence-based DL architectures have demonstrated a strong capacity to extract contextual representations that generalise well to peptide developability tasks [Guntuboina et al., 2023, Zhang et al., 2026]. Co-folding representations offer a complementary axis: by capturing target-binder relationships such as spatial proximity and physicochemical complementarity, they encode information that one-dimensional sequences obscure [Jumper et al., 2021, Abramson et al., 2024, Passaro et al., 2025]. However, the interplay between protein language and protein folding models has not been extensively explored for peptides, despite showing promising outcomes for full-length globular proteins [Qurat-ul-ain et al., 2026, Li and Luo, 2025]. Moreover, existing comparisons across these representation types remain fragmented: new architectures are typically benchmarked against prior specialised models rather than against strong classical baselines, embeddings are evaluated off-the-shelf rather than fine-tuned or fused across representation types, and co-folding-derived affinity prediction has yet to be adapted or assessed for peptide-length binders at all. To close this gap, we present a critical assessment of chemical descriptors, PLM and folding model representations for predicting therapeutic peptide developability and target binding affinity: we benchmark these representations against classical physicochemical baselines, systematically fine-tune and fuse embeddings across representation types, and adapt Boltz-2’s affinity module to peptides, assessing it alongside the other methods. Together, this offers a robust, unbiased evaluation of current fundamental methods for peptide scoring and filtering to aid candidate selection in *de novo* generative pipelines.

## 2 Results

### 2.1 Assembling a robust dataset benchmark for peptide scoring tasks

To establish a robust benchmark for therapeutic peptide developability, we first collected and harmonised five diverse datasets spanning aggregation, membrane permeability, non-fouling, kinetic stability, and thermodynamic stability (Table 1). These properties were selected to cover a large space of traits to optimise in therapeutic designs [Wang et al., 2022, Guntuboina et al., 2023]. The reliability of predictive models depends heavily on the quality of their training, validation, and test splits. As shown in Figure S1, our leakage-aware splitting strategy (see Section 5.2 for details) uniformly reduces peptide sequence leakage across the benchmarked datasets when compared to the original splits provided in the source publications.

In addition to these developability traits, we curated a binding affinity dataset by aggregating four sources, yielding over 8,500 entries (Table 1). To our knowledge, this is the largest peptide binding affinity datasets curated to date. Partitioning this dataset requires a different approach, because each measurement is attached to two sequences rather than one. Existing small molecule affinity models control only the target side, excluding training proteins above a sequence identity threshold to the benchmark [Passaro et al., 2025, Shenoy et al., 2026]. Mattsson and Walters [2026] have shown this to be insufficient: homologous proteins below the threshold retain highly correlated binding profiles, and a ligand-only baseline with no protein information reaches r = 0.66 on the FEP+ benchmark ([Wang et al., 2015]), indicating that much of the reported performance reflects memorisation of the ligand rather than learned biophysics. We therefore developed a graph-based partitioning procedure (Section 5.2) that constrains similarity on both sides of the pair simultaneously, so that no test complex shares a similar peptide or a similar target with anything seen in training. By construction, this strategy removes entirely the data leakage issue based on sequence homology (Figure S1).

**Table 1:**
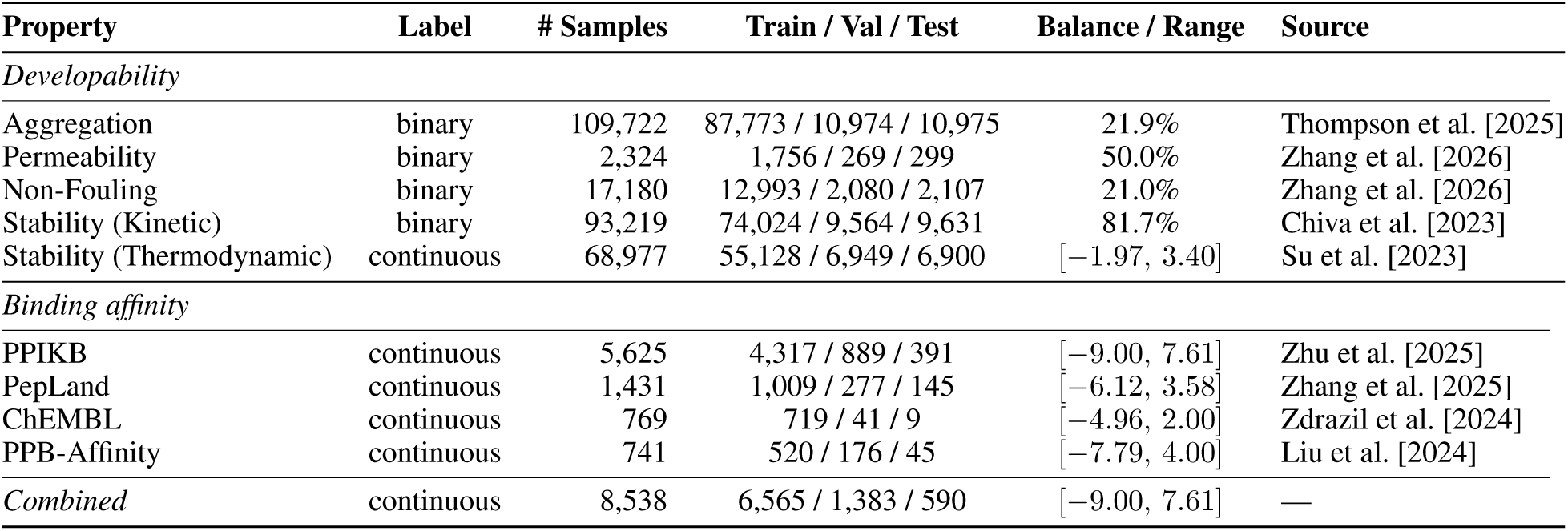
Summary of datasets used. *Balance / Range* reports the positive-class fraction for binary tasks and the label value range for continuous tasks. Binding-affinity labels are log_10_of the measured affinity in µM; the merged set combines *K_D_* (58.8%), IC_50_(29.3%), *K_i_* (9.3%) and other measurement types (2.6%).

### 2.2 Peptide chemical descriptors convey essential information to predict developability properties

Physicochemical features have provided a valuable toolkit in combination with machine learning to identify high-level protein functional traits [Ofer and Linial, 2015]. To explore the usefulness of sequence-derived chemical descriptors in predictive peptide tasks, we computed 45 descriptors spanning charge and electrostatics, hydropathy, size and composition, flexibility, secondary structure propensity, hydrogen bond capacity, disorder and degron signals, and post-translational modification motifs (Figure S2, see Section 5 for further details).

Taken individually, these descriptors are highly informative for some tasks and close to uninformative for others (Figure S2). Charge dominates permeability, with net charge and isoelectric point alone reaching a single-feature ROC-AUC of 0.85, while hydrophobicity, terminal flexibility and peptide size drive non-fouling. For thermodynamic stability, hydrophobicity shows the strongest monotonic trend, though a modest one (Spearman ρ ↑ 0.25). By contrast, no single descriptor separates the aggregation or kinetic stability classes appreciably better than chance.

We therefore asked whether the descriptors are jointly more informative than any of them in isolation. Projecting each dataset onto its first two supervised PLS latent variables (Figure 1) recovers a clear separation for permeability (out-of-fold AUROC 0.86) and non-fouling (0.82), but only a weak one for aggregation (0.64), kinetic stability (0.65) and thermodynamic stability Spearman ρ = 0.38). Because PLS is linear, these values are a conservative lower bound. The permeability result reflects a real data limitation: the positive class consists of cell-penetrating peptides, a fraction of which are engineered arginine-rich constructs (Figure S3). The non-fouling dataset carries a milder confound of a different kind, in that its non-fouling class contains only short peptides (5-11 residues) whereas its fouling class extends to longer sequences. Restricting the analysis to a length-matched, class-balanced subset (↓ 11 residues, n = 7,198) reduces the separation from AUROC 0.82 to 0.74 but does not remove it, and the projection is unchanged when every size-dependent descriptor is excluded. The signal is instead carried by hydrophilicity, low ε-sheet propensity and disorder-promoting composition, consistent with the established biophysical basis of fouling resistance (Figure S2) [Nowinski et al., 2012].

Given this separability, despite weak for several tasks, we asked whether a gradient-boosted tree model (XGBoost) ([Chen and Guestrin, 2016]) could recover more of the signal at negligible computational cost. The results showed improvement on every task (Aggregation: AUROC = 0.64 → 0.70; Permeability: AUROC = 0.86 → 0.93; Non-fouling: AUROC = 0.82 → 0.84; Kinetic stability: AUROC = 0.65 → 0.73; Thermodynamic stability: Spearman *ρ* = 0.38 → 0.47, Figure 2).

Overall, chemistry alone accounts for the tasks whose determinants are essentially compositional, but leaves most of the signal in the aggregation and stability tasks unexplained. We therefore asked next whether learned representations from protein language and folding models can fill this gap.

**Figure 1:**
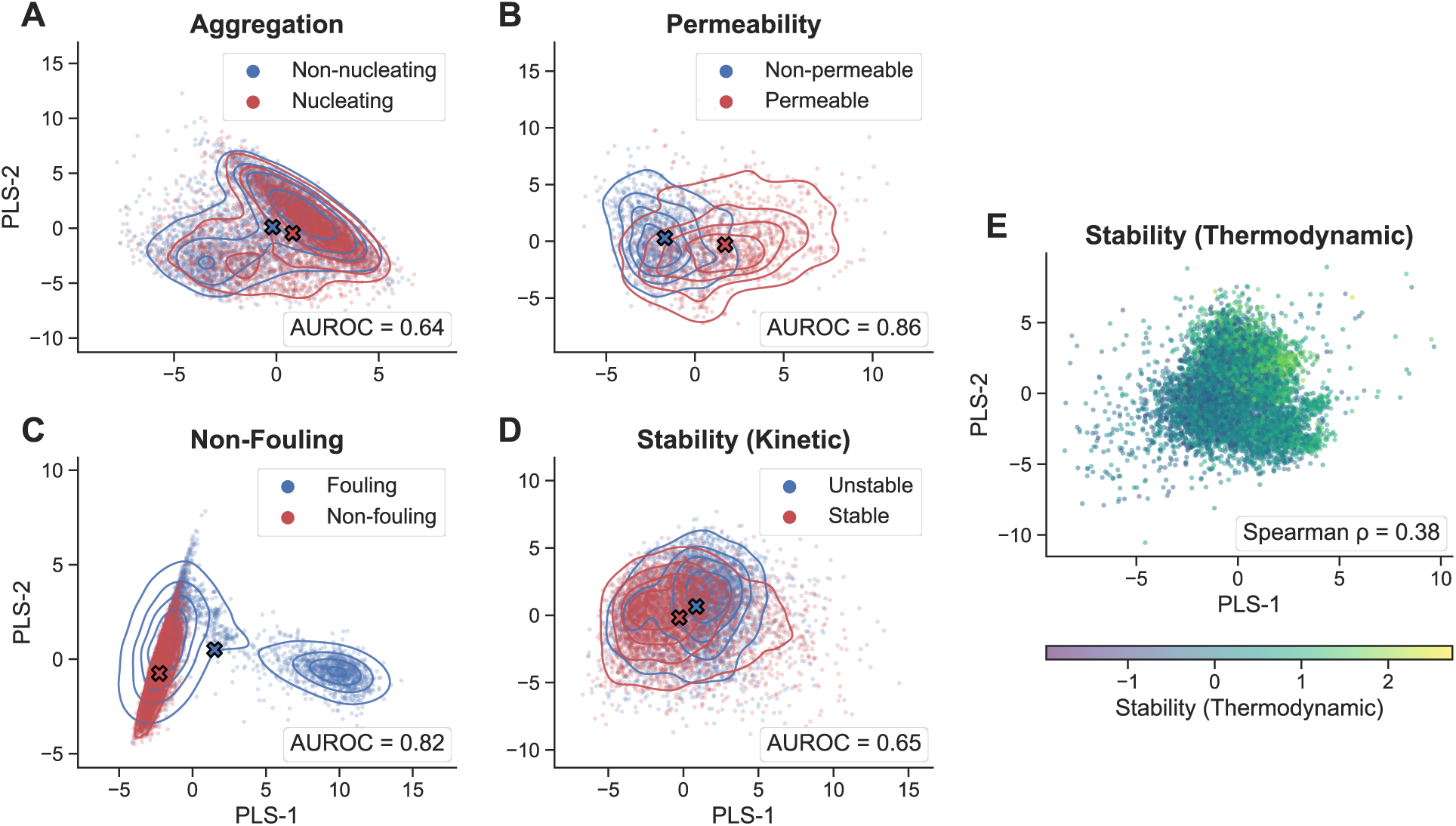
Separation of developability classes by peptide chemistry, across the five benchmark tasks. Each panel is a two-dimensional partial least squares (PLS) projection of per-peptide physicochemical properties. (A-D) Binary classification tasks, coloured by class. Solid lines are per-class kernel-density contours and the crosses mark the class centroids. (E) Thermodynamic stability (SaProt), a regression task, with points coloured by the continuous stability score. Points are randomly subsampled for display, whereas the PLS projection is fit on the full dataset. Cross-validated linear separability is assessed using AUROC for classification tasks and Spearman correlation for regression.

### 2.3 ESM2 and Boltz-2 exhibit similar performance, improved by task-dependent fine-tuning

Evolutionary Scale Models (ESM2, Lin et al., 2023) have proven valuable across predictive tasks, including multi-label classification of antimicrobial peptides [Bin et al., 2025], detection of hydrophobic patches on protein surfaces [Gogishvili et al., 2024] and assessment of folded-conformer stability [Li and Luo, 2025]. We derived ESM2 embeddings for each curated dataset and attached an MLP head to predict the task label (binary classification, except for Thermodynamic Stability, a regression task; see Section 5 for details). With frozen embeddings, ESM2 does not surpass the XGBoost baseline trained on the chemical space of peptides on any task except Thermodynamic Stability, where Spearman ρ rises from 0.47 to 0.58 (Figure 2). Fine-tuning recovers much of the remaining gap on the stability and aggregation datasets, but leaves permeability and non-fouling essentially unchanged, suggesting that performance on these two tasks is already saturated.

We next asked whether the pair-representation embeddings of a DL folding model also carry information relevant to peptide developability. Our initial expectation was that predicted structural features would be less reliable for peptides than for longer proteins, given the intrinsic flexibility that makes peptides hard to crystallise, and which is why isolated peptides are more often characterised by NMR than by X-ray crystallography. Contrary to this hypothesis, the pLDDT and confidence scores of Boltz-2 generated structures are high across datasets (Figure S4C). Only the Kinetic Stability dataset shows lowered confidence, consistent with its enrichment in very short peptides (median length 13 residues, with 28% of sequences at 10 residues or fewer, against 43-50 residues for the Thermodynamic Stability set). In line with this, frozen embeddings from the Boltz-2 trunk match or exceed the performance of frozen ESM2 embeddings on the predictive benchmark. While fine-tuning provides a substantial performance boost for ESM2, fine-tuning the Boltz-2 trunk yields only marginal additional gains (Figure 2).

Interestingly, an inverse folding model such as ProteinMPNN, which operates on the same predicted structures, underperforms. To understand why, we compared the perplexity of the native peptide sequences with that of randomly scrambled controls threaded onto the same backbone (Figure S4A). The native sequence scores below its scramble for 89-94% of peptides in the four short sets and for 99.8% of the thermodynamic stability set, confirming that ProteinMPNN is able to extract structural signal. However, the gap is narrow and sequence-length-dependent, being the largest for a positive control of 24 small single-domain natural proteins (between 53 and 126 residues) taken from the two-state folding benchmark of Jackson [1998]. Native perplexity falls with length (Figure S4D), and for the shortest peptides it approaches the uninformative limit of 20 expected under uniform guessing over the amino acid alphabet. Length dependence is not specific to inverse folding: ESM2 pseudo-perplexity follows the same trend (Figure S4D) and leaves an even smaller native-versus-scrambled gap on the short peptides (Figure S4B). Conditioning on the predicted backbone therefore does add information beyond sequence, since ProteinMPNN scores the native sequence mostly below ESM2, and the folds themselves are not the bottleneck: Boltz-2 reports them as confident throughout (Figure S4C). Possibly, what limits ProteinMPNN is the amount of context available from small peptides and by its comparatively shallow architecture relative to ESM2 and Boltz-2 [Dauparas et al., 2022].

**Figure 2:**
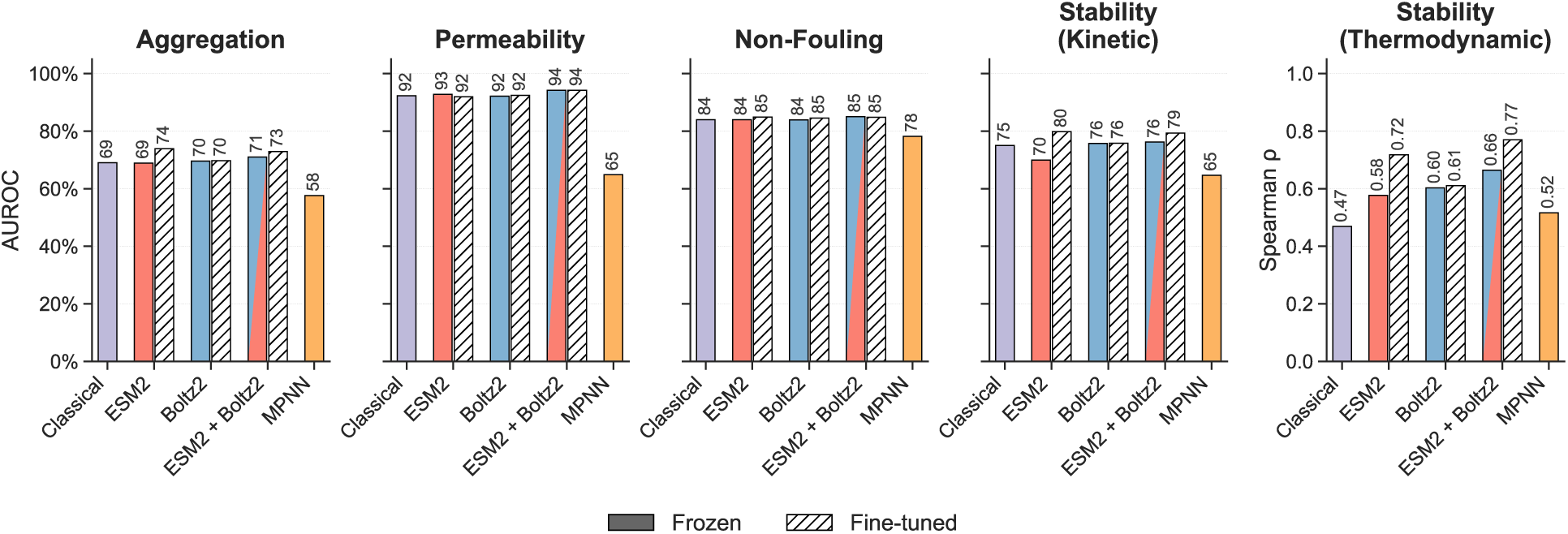
Classical machine learning versus DL baselines across five peptide-property tasks. Each panel compares a classical XGBoost head on physicochemical descriptors (“Classical”) with four DL architectures (ESM2, Boltz-2, fused ESM2-Boltz-2, ProteinMPNN; model embeddings + MLP head). DL baselines were obtained using either frozen (solid) or fine-tuned model weights (hatched). Performance is AUROC for the four classification tasks and Spearman ρ for thermodynamic stability (regression, own axis); higher is better. Fusion results are shown for concatenated embeddings (see Section 5.3.5 for more details)

### 2.4 Fusing model embeddings matches or overtakes unimodal baselines

Given the strong unimodal ESM2 baselines and the largely matching contribution of structural inputs, we asked whether the geometric and pairwise information encoded by the Boltz-2 trunk could provide complementary signal when explicitly fused with the sequence backbone.

We compared three fusion strategies. All three project the ESM2 embedding, the Boltz-2 single representation and the row-averaged Boltz-2 pair representation to a common width at each residue, and differ only in how the three are combined: sum adds them element-wise, concatenation stacks them along the channel axis, and gating modulates each by a learned sigmoid gate conditioned on all three, plus a residual term (Equation (3)). The fused features are mean-pooled and passed to a shared MLP head. Each strategy was evaluated with ESM2 either frozen or fully fine-tuned, with Boltz-2 held frozen throughout due to computational constraints (Figure 2 and Table S1). When both encoders are frozen, the best fusion variant improves on the better of the two unimodal encoders across all five tasks, though by modest margins: from 69.6 to 71.4 AUROC on aggregation, 92.8 to 94.4 on permeability, 84.0 to 85.0 on non-fouling, and 75.7 to 76.7 on kinetic stability. The exception in magnitude is thermodynamic stability, where Spearman ρ rises from 0.603 to 0.684, a gain roughly five times larger in relative terms than any of the classification tasks. This is also the only setting in which the two encoders are clearly contributing different information rather than the same sequence prior.

Fine-tuning ESM2 largely absorbs this advantage. On aggregation and kinetic stability the best fused model does not improve on the fully fine-tuned unimodal ESM2 baseline. Only two tasks retain a benefit from fusion once the sequence encoder is trainable: thermodynamic stability, where ρ improves from 0.717 to 0.770, and permeability, where AUROC improves from 91.9 to 94.8. The permeability result is notable in that full fine-tuning alone marginally degrades performance relative to the frozen baseline (91.9 against 92.8), which the structural input then more than recovers. The overall pattern therefore mirrors the one observed for the unimodal baselines: tasks that are already near saturation, such as permeability and non-fouling, absorb additional representational capacity poorly, whereas thermodynamic stability, the only task with substantial headroom, benefits from every increase in capacity we applied. Differences between the three fusion architectures are small, and their ranking depends on the ESM2 regime rather than being consistent across it. We note that Boltz-2 remained frozen in all fused configurations for compute reasons, so these results characterise fusion onto a fixed structural representation and may understate what joint fine-tuning of both encoders could achieve.

### 2.5 Binding affinity

Given the predictive strength of Boltz-2 trunk outputs and ESM2 embeddings on peptide-only developability tasks, we explored their capacity to predict protein-peptide binding affinities. While leveraging such representations for binding affinity has been attempted for protein-protein interactions, albeit with limited success [King et al., 2025, Alsamkary et al., 2025, Liu et al., 2024], their application to peptide-protein interactions remains largely unexplored. To address this, we systematically evaluated these representations across distinct architectures. We first established a baseline by pooling the representations into one-dimensional vectors and passing them through simple MLPs. Next, we investigated the Boltz-2 affinity module (AM) [Passaro et al., 2025] under two conditions: a zero-shot evaluation to determine if its pre-trained weights (originally fit on small molecules) carry intrinsic predictive power for peptides, and a fully trained module fine-tuned explicitly on our peptide dataset.

Evaluation was performed on 14 held-out panels (Figure 3) from the test set. Each a set of measurements (>10 data points) comes from a unique publication and shares a binding label type (i.e. IC50) and target sequence. As a trivial baseline, we included the correlation of peptide molecular weight (MW) with the labels. Averaged over the 14 panels (Figure 3), the Boltz-2 pair representation performs best (mean ρ = 0.44), with ESM2 embeddings (0.40) trailing not far behind. All three correlate positively on the large majority of panels, though each fails outright on a few. The pre-trained affinity module applied zero-shot perfoms poorly across all panels (ρ = 0.07). Training the AM on our curated peptide binding affinity data improves relative to its zero-shot performance by Δ*ρ* = 0.28.

In classical drug discovery, the maximal affinity of ligands is known to correlate strongly with the number of non-hydrogen heavy atoms [Kuntz et al., 1999]. The Boltz-2 authors have introduced a correction to account for this bias [Passaro et al., 2025] in their affinity module. The same relationship holds in our curated peptide dataset (Figure S5)A: MW alone correlates with measured affinity at *ρ* = →0.32 over all peptides in the dataset, heavier binders binding more tightly. Our model learns this relationship and overstates it. Its predictions correlate with MW at *ρ* = →0.72 across the same datapoints (Figure S5B). It reflects a fundamental biophysical reality, i.e. larger molecular surfaces inherently increase baseline van der Waals and hydrophobic interactions, rather than an artefact of the computational embeddings. Ranking peptides by weight alone reaches a mean *ρ* of 0.35 across the 14 panels, ending best method on five panels (Figure 3).

We next examine the strongest of the four models, the Boltz-2 pair representation, panel by panel (Figure 4). Its predictions occupy a much narrower range than the measurements in every one of the 14 examined assays: within a panel they span a mean standard deviation of 0.12 log units against 0.93 for the measured labels, and every panel is centred between 3 and 20 µM, regardless of whether its measured labels fall in the nM or mM range. This behaviour likely reflects limitations observed during model training. The model fits the training split readily but the validation loss spikes after five epochs. Early stopping preserves the ranking signal, whereas later checkpoints display a wider predicted range with poor ranking signal. This behaviour reflects the difficulty of the task under our partitioning scheme. Training data occupies a single connected component of the similarity graph (Section 5.2), so every validation and test complex differs from anything seen in training on both the peptide and the target side, making generalisation tougher. The narrow predicted range suggests the model retreats towards the training mean rather than extrapolating. Training difficulties due to data limitations extend to other models as well, so this behaviour is not limited to a particular architecture.

The consequence of the narrow range of predicted affinity labels is that ranking and absolute accuracy come apart entirely: the model can order candidates against a target, within the same type of assay, but it cannot identify a nanomolar binder as one. This distinction is recognised in the field, separating hit discovery, which requires distinguishing binders from non-binders across a library, from hit-to-lead optimisation, which requires resolving fine-grained differences among known binders, and supervise the two with separate heads (Passaro et al. [2025], Zhu et al. [2013]). Structure confidence does not have a significant impact on the predictions: a panel’s mean iPTM is uncorrelated with how well its peptides are ranked (Figure 4, *ρ* = −0.14).

**Figure 3:**
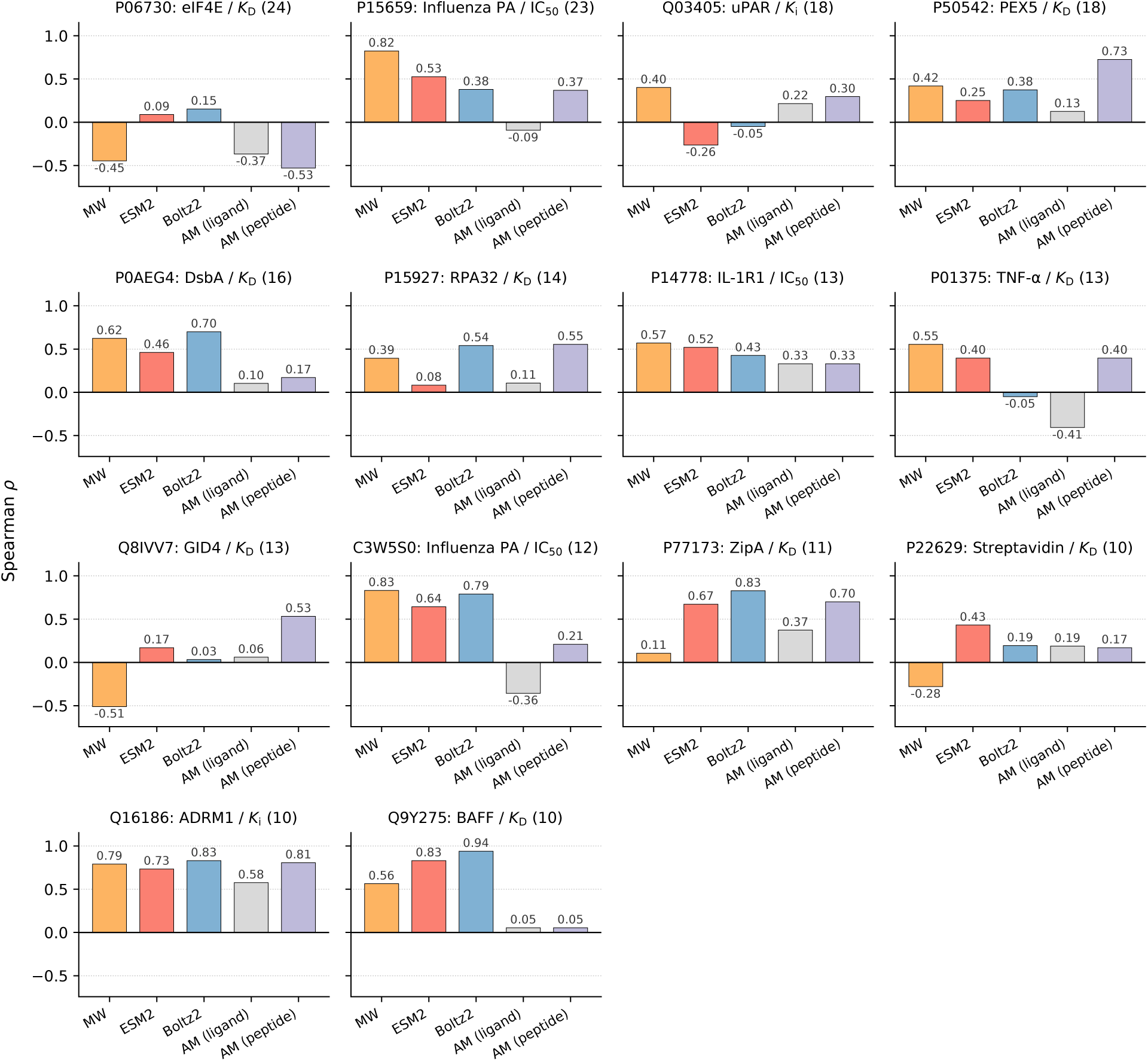
Spearman *ρ* between predicted and measured binding affinity for the fourteen groups of the evaluation set. Each panel title is structured as following: UniProt ID: Protein name / Assay label (number of measurements). Bars, left to right: MW, sign-inverted so that a higher correlation means heavier peptides bind more tightly; ESM2 sequence embeddings + MLP; Boltz-2 pair representation embeddings + MLP; AM (ligand), Boltz-2 built-in affinity module and AM (peptide), the same module architecture trained on our peptide dataset.

**Figure 4:**
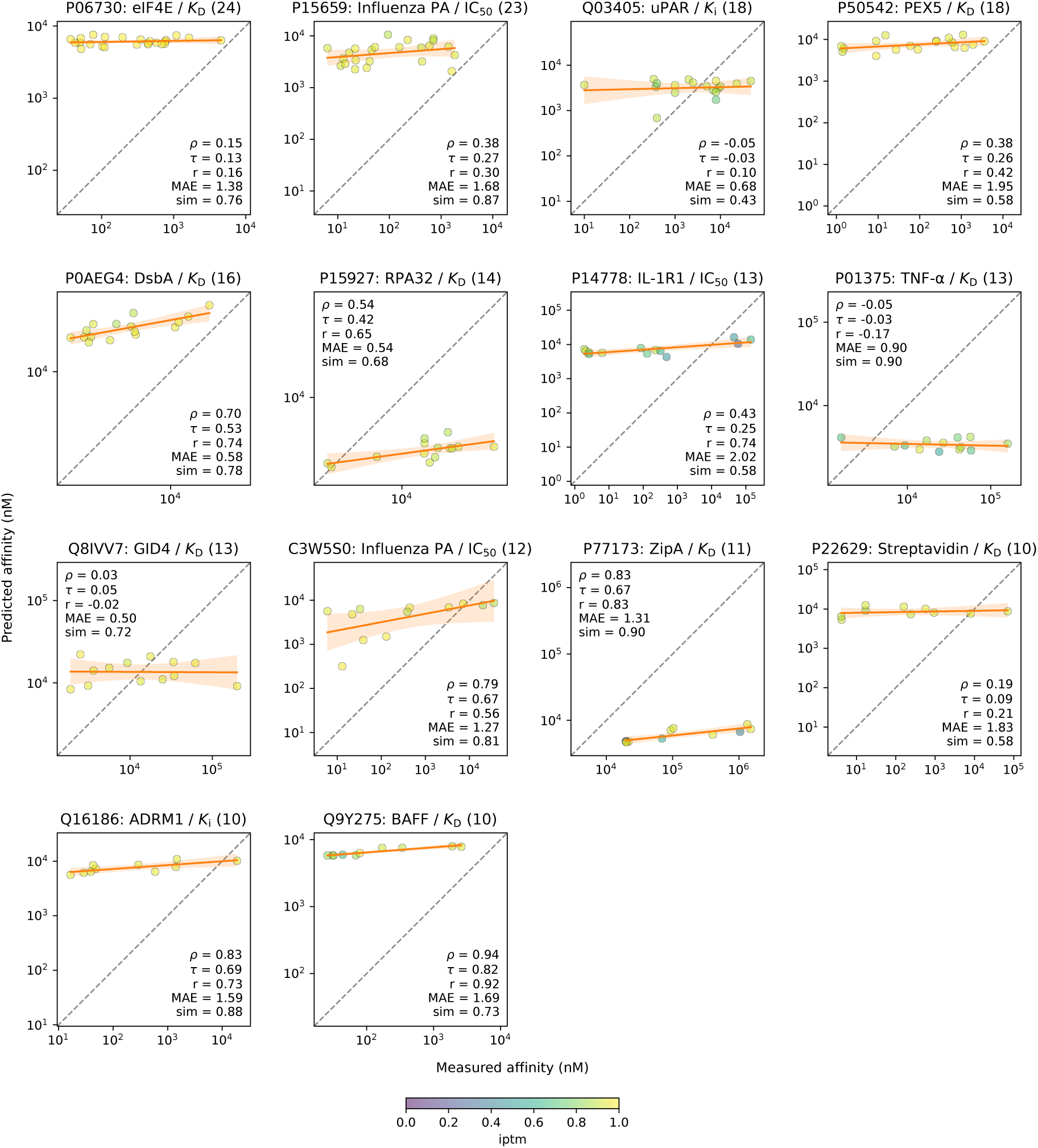
Predictions from the pair representation of the pretrained Boltz-2 model, passed to an MLP head. Panels are the the fourteen groups of the evaluation set. The orange line is the least-squares fit with a 95% confidence band. Points are coloured by the interface pTM (iPTM) of the Boltz-2 complex prediction. Each panel reports Spearman **ρ**, Kendall *τ*, Pearson *r*, mean absolute error (MAE), and *sim*, the mean pairwise Levenshtein ratio between the distinct peptides in the panel. Each panel title is structured as following: UniProt ID: Protein name / Assay label (number of measurements)

## 3 Discussion

Across a range of peptide developability tasks, classical models trained on physicochemical descriptors matched or approached DL architectures at a fraction of the computational cost (Table S2). ESM2 and Boltz-2 improved on them only marginally, and fine-tuning either model carried a further limited advantage over frozen weights while adding substantial cost, particularly for the Boltz-2 trunk. Similarly, fusing these embeddings yielded small performance gains despite a significant increase in model complexity. The explanation likely lies in the nature of the representations. If a developability metric is primarily driven by a few basic physicochemical descriptors, a DL model that simply learns to extract these same traits from the sequence will inherently offer little additional predictive power. Membrane translocation is governed by charge and lipophilicity, resistance to fouling is driven by hydrophilicity, low ε-sheet propensity and disorder-promoting composition. This is not limited to peptides. Ofer and Linial [2015] showed a decade ago that sequence-derived biophysical features paired with standard classifiers match dedicated predictors on nucleic-acid-binding tasks without alignment or evolutionary information, and that feature selection reduces the set to fifteen interpretable descriptors with almost no loss. Deng et al. [2023] report the same pattern for small molecules, with random forests on RDKit descriptors outperforming SMILES- and graph-based representation learning across most property prediction benchmarks. Beyond cost, descriptors offer interpretability that pooled embeddings cannot. They identify which quantities drive a prediction, which is directly actionable in a design context, whereas an embedding gives a number without an account of what produced it. It is worth noting that the co-folding models themselves internalise physicochemical features. Boltz-2 inherits AlphaFold3’s input featurisation, in which every residue and ligand is embedded from an RDKit reference conformer and each of its atoms carries its element identity, its formal charge and its reference coordinates as explicit input features [Abramson et al., 2024, Passaro et al., 2025]. Charge and local geometry, two of the descriptor families we compute, are therefore supplied to the network rather than learned from scratch, albeit at atomic resolution rather than as whole-molecule summaries.

Beyond developability trait predictions, our results indicate that DL representations carry usable signal for ranking peptide binders against a protein target, with the Boltz-2 model being the most predictive. However, the narrow range of predicted affinities means the model orders candidates within an assay without resolving their absolute potency. Nonetheless the ordering signal is accurate, and we expect performance to improve with more data, as demonstrated for small molecule affinity modules trained on millions of measurements [Passaro et al., 2025, Shenoy et al., 2026]. On the contrary, the Boltz-2 AM applied zero-shot has no predictive capacity for peptides, which we attribute to tokenisation differences: small-molecule binders enter Boltz-2 as SMILES and are tokenised per atom, whereas a peptide supplied as a protein chain is tokenised per residue, so the module receives inputs of a kind it was never trained on. That the same architecture recovers usable performance once trained on peptide data suggests the limitation lies in the pre-training distribution rather than in the architecture itself. MW ranks peptides nearly as well as any model we tested. This is a long-recognised confound in drug design [Kuntz et al., 1999]: larger molecules present more surface for contact, and binding strength rises with it. A model that learns this dependence is not thereby intrinsically wrong, since the trend is genuine. This, however, raises design limitations, because the same bulk that improves binding degrades other traits: molecules of high MW and lipophilicity tend to violate Lipinski’s rules, showing reduced permeability, poorer solubility and faster hepatic clearance [Hopkins et al., 2004]. A more fundamental limitation concerns what the pair representation encodes. Co-folding models derive it by mining coevolutionary signal from multiple sequence alignments, which presupposes that the sequences in question have an evolutionary history. Designed peptides do not: they are synthesised in vitro, often from randomised libraries, and are not sampled from any evolutionary space. The signal on which the representation is founded is therefore weak or absent on the binder side of the complex, present only for the target. This compounds a broader observation that co-folding models capture interaction geometry without learning the underlying physics [Masters et al., 2025]. Changes in binding potency frequently turn on subtle stereochemical or chemical distinctions (e.g. two near-identical peptides differing only in the chirality of a residue can differ entirely in affinity) and a representation built from sequence-level covariation has no obvious mechanism for capturing them.

## 4 Conclusions

By comprehensively assessing sequence- and structure-derived features across five peptide developability tasks, as well as predicting their binding affinity, we have shown that:

- Inexpensive, explainable physicochemical descriptors paired with classical machine learning (like XGBoost) match or approach the performance of complex DL architectures across multiple developability tasks. They achieve this at a drastically lower computational cost while maintaining high interpretability.
- Fusing protein language and folding model representations provides only marginal performance gains over unimodal baselines. When tasks are already near saturation, they absorb additional DL capacity poorly.
- While the Boltz-2 pair representation effectively ranks peptide binders against a target, its absolute potency predictions are heavily biased by MW. The model tends to assume heavier peptides bind more tightly, which actively conflicts with medicinal chemistry goals of minimizing bulk to preserve solubility and permeability.
- With fewer than 0.2% of Protein Data Bank (PDB) entries annotated as peptides [Park et al., 2024], the performance of all evaluated DL models is strictly bounded by a lack of structural coverage and biochemical measurements

Taken together, these results suggest that near-term progress on peptide developability and binding affinity predictions will come less from larger models than from broader and better-annotated peptide data. Interpretable descriptor baselines should be reported alongside DL models as a check on where the added capacity is genuinely earning its place.

## 5 Materials and Methods

### 5.1 Data curation

#### 5.1.1 Developability data

Kinetic (proteolytic) stability data were taken from Chiva et al. [2023], in which tryptic peptides are assigned to degradation clusters from time-course mass-spectrometry intensities: the ambiguous cluster was discarded, the remaining two clusters were mapped to unstable (0) and stable (1), and peptides with more than three missing time points were removed. Thermodynamic stability data were taken from the proteolysis-based stability scores of Rocklin et al. [2017] as redistributed in the SaProt benchmark [Su et al., 2023]; replicate measurements of the same sequence were collapsed to their mean. Aggregation data were taken from the CaNYA random-peptide amyloid nucleation assay [Thompson et al., 2025] (GEO accession GSE268261, NNK1-4 libraries plus an independent replication library); sequences were truncated at the first stop codon and discarded if shorter than five residues, duplicate sequences were resolved in favour of the replication library and then the held-out library, and the binary nucleator call was used as the label. Non-fouling and cell-permeability data were taken from PeptiVerse [Zhang et al., 2026] and deduplicated on sequence; the permeability positives are cell-penetrating peptides drawn from 22 independent studies and the negatives are UniProt-derived peptides sampled to match the positive length distribution. All datasets were mapped onto a single schema (peptide identifier, sequence, label, label units, label type, original publication split and our computed split), with one row per unique peptide and a deterministic identifier.

#### 5.1.2 Binding affinity data

Binding affinity data were assembled from four sources and expressed on a common log_10_(µM) scale. From ChEMBL v36 [Zdrazil et al., 2024], we retrieved every molecule annotated `molecule_type = Protein` together with its HELM notation [Zhang et al., 2012] and canonical SMILES, reconstructed the one-letter peptide sequence from HELM, and joined all Ki, Kd, IC50 and EC50 activities to their assay, target and document records, keeping only single-component targets so that each measurement maps to one UniProt sequence. Following Landrum and Riniker [2024], we retained only activities with no data-validity comment, an exact standard relation, a valid pChEMBL value and a positive nanomolar concentration, a peptide of at least two residues with an available SMILES string, a biochemical assay type, a named single-protein target with confidence score 9, and an assay description containing no mutant, mutation or variant annotation. Protein-peptide interactions were retrieved from the Protein-Peptide Interaction Knowledge Base (PPIKB) [Zhu et al., 2025], from which we extracted the target and peptide sequences, the reported affinity (Kd, Ki or IC50) and the PubMed identifier of the source publication. Cyclic peptides were removed, two errors identified against the primary literature were corrected (three HLA-DRB1 alleles collapsed into a single target sequence for PMID 29317506, and fabricated IC50 records together with mis-transcribed Kd/Ki values for PMID 34589387), allele-specific entries were dropped, and a SMILES string was generated for every remaining peptide, via the Pistoia HELM Core Library ^1^. We recovered each PubMed identifier from the RCSB REST API, so that measurements could be attributed to their source publication. In addition, we incorporated PPB-Affinity [Liu et al., 2024], which aggregates protein-protein binding affinities from SKEMPI v2.0, AB-Bind, SAbDab, PDBbind, Affinity Benchmark and ATLAS, each entry annotated with a PDB code and its receptor and ligand chains. Peptide-protein affinities were additionally taken from the canonical-residue binding subset distributed with PepLand [Zhang et al., 2025], from which we retained the target sequence, the peptide sequence, the PDB identifier of the complex and the measurement kind (Kd, Ki or IC50). PepLand reports affinities as pKD, which we mapped onto the common scale as log_10_(µM) = 6 - pKD, and we recovered the PubMed identifier of each entry from its PDB accession through the RCSB REST API so that measurements could be attributed to a source publication.

Because public assay collections carry heterogeneous and often incomplete metadata, we treated affinity and potency labels (Kd, Ki, IC50, EC50) interchangeably. Duplicated and inconsistent measurements were then removed in two stages. ChEMBL and PPIKB records were pooled and keyed by (SMILES, target sequence, measurement type) so that paper-level disagreement is estimated across both sources simultaneously: where both sources reported the same interaction from the same publication the ChEMBL record was kept, publications in which more than 60% of at least five overlapping pairwise comparisons disagreed by more than one log unit were flagged, duplicate measurements differing by more than tenfold were reduced to the record from the best-supported publication, and groups containing either exact duplicates or disagreements of at least three log units were collapsed to the measurement closest to the group median. The pooled set was then concatenated with PepLand and filtered once: duplicates were collapsed across sources by (target, ligand, measurement type), preferring records that report an assay method and averaging the survivors; only complexes containing at least one chain of 50 residues or fewer were retained; entries containing non-canonical residues were dropped; each target was capped at 15 measurements, homogenously sampling the affinity distribution, to reduce oversampling for large clusters; and complexes exceeding 1,050 residues in total were removed. Multi-chain targets were excluded from the final dataset.

### 5.2 Dataset partitioning

#### 5.2.1 Sequence-based partitioning

Partitioning of the developability datasets was performed using a strict implementation of the Fast-Part algorithm [Ahmed et al., 2024] to prevent data leakage between training, validation, and test sets. Sequences were first deduplicated, then clustered using MMseqs2 [Steinegger and Söding, 2017] at 30% sequence identity with 80% alignment coverage. An initial draft partition was constructed by greedily assigning entire clusters to validation (10%) and test (10%) sets, with the remainder allocated to training (80%). A leakage audit was then performed using DIAMOND [Buchfink et al., 2021] to identify training sequences with >30% similarity (for the validation-test audit for SaProt dataset a threshold of >50% was used due to high sequence similarities) to the validation or test sets; any such sequences were reassigned to the corresponding evaluation set to eliminate cross-partition homology. Finally, split labels were propagated to all duplicate sequences sharing the same canonicalised representation. This procedure was applied independently to each dataset to produce a new train/validation/test split.

In addition, for datasets that underwent splitting in previous publications, we kept the paper splits in order to analyse leakage compared to our splits.

#### 5.2.2 Graph-based partitioning

For the binding affinity dataset, all unique chains were extracted from the peptide and target sequences of every measurement. Homologous chain pairs were identified by an all-versus-all DIAMOND search [Buchfink et al., 2021] at 30% identity (more-sensitive mode, no cap on hits per query). Because DIAMOND’s seed-and-extend heuristic rarely reaches significance on short peptides (fewer than 1% of chains below 11 residues receive any hit) all chains of at most 20 residues were additionally compared exhaustively and linked where length-normalised Levenshtein similarity reached 0.8.

We then built one undirected graph over measurements and chains, joining each measurement to its constituent chains and each chain to every chain found similar by either search. Its connected components, which we term islands, are closed under similarity by construction, so no chain in one island is similar to a chain in another. Hence, we can assign these islands to splits leakage free. To evaluate model performance under realistic experimental constraints, we group these measurements into *panels*. We define a panel as a set of measurements sharing a single source publication, measurement type, and target sequence (i.e., one `pubmed_id::label_type::target_sequence` group). Because rank correlations over small sample sizes are dominated by noise, we only consider panels containing at least ten measurements as viable for evaluation.

**Table 2:** Island assignment from graph-based partitioning. A panel represents one publication’s measurements of one assay against one protein. The final column counts panels holding at least ten measurements, the threshold required for robust per-assay rank correlation. No island spans more than one split, so the island counts are additive.

| | Islands | Labels | Panels | Panels $\geq 10$ |
| --- | --- | --- | --- | --- |
| Train | 1 | 6,565 | 3,225 | 82 |
| Validation | 229 | 1,383 | 787 | 0 |
| Test | 221 | 590 | 352 | 14 |
| Total | 451 | 8,538 | 4,364 | 96 |

The similarity graph is dominated by a single component holding 76.9% of the points in the dataset; it cannot be subdivided without breaking the leakage guarantee above, and so becomes the training set. Of the remaining 450 islands, we assign every island containing an evaluable panel (at least ten measurements) to the test set and the rest to validation (Table 2). This results in validation having 1,383 measurements to drive the loss and early stopping. Models are evaluated within assay panels rather than over the held-out test split as a whole. We retain the 14 largest, manually corrected assay panels, each holding at least ten measurements, 211 measurements in total.

### 5.3 Feature extraction and representations

#### 5.3.1 Physicochemical descriptors

Forty-five descriptors were computed for each peptide: thirty-four from the amino-acid sequence and eleven from a molecular graph derived from it. Net charge at pH 7.4, isoelectric point, average MW, GRAVY on the Kyte–Doolittle scale, aromaticity, the instability index and the helix, turn and sheet prevalence ratios were obtained using Biopython v1.84 [Cock et al., 2009]; the charge distribution asymmetry *k*, the charge–proline distribution asymmetry Ω, the polyproline-II propensity and counts of charged, positive, negative, neutral, expanding and disorder-promoting residues and of phosphorylatable sites were taken from `localCIDER v0.1.21` [Holehouse et al., 2017]. We additionally recorded sequence length, the cysteine count, the summed Pro, Glu, Ser and Thr content, the numbers of proline-directed phosphorylation sites ([ST]P) and of C2H2 zinc-finger motifs, and the largest hydropathy difference between any two five-residue windows. The remaining nine sequence descriptors reduce three per-residue profiles (i.e. hydropathy, charge at pH 7.4 and backbone flexibility [Vihinen et al., 1994]) by four position-aware operators: the profile mean, a right-weighted mean that is positive when large values accumulate towards the C-terminus, a termini-weighted mean that up-weights both ends relative to the centre, and a burden that sums only the positive part of the profile. Flexibility is defined by a nine-residue window and is therefore undefined for peptides of nine residues or fewer (16% of curated peptides), for which all three flexibility descriptors were set to missing. The remaining eleven descriptors were computed with RDKit, the open-source cheminformatics package ^2^, as exposed by `cyclicpeptide v1.4.2` [Yang et al., 2025], from a linear neutral SMILES generated from the canonical sequence with p2smi: monoisotopic mass, topological polar surface area, logP, molar refractivity, heavy-atom count, hydrogen-bond donor and acceptor counts, rotatable bonds, aromatic rings, the fraction of sp^3^-hybridised carbons, and Morgan fingerprint density at radius one. Descriptors constant within a dataset were dropped before training, leaving between forty-three and forty-five descriptors per dataset. Any missing values (such as flexibility scores for short peptides) were imputed using the median of that feature’s distribution in the training split.

#### 5.3.2 Protein language models

ESM2 [Lin et al., 2023] served as our sequence encoder, in its 650M-parameter configuration (`esm2_t33_650M_UR50D`; 33 transformer layers, 1280-dimensional representations). For embedding extraction, sequences were grouped into batches under a budget of 4096 tokens rather than a fixed number of sequences, so that memory use does not depend on the length distribution of the dataset. We retained the per-residue representations of the final transformer layer, excluding the classification and end-of-sequence tokens, giving one L x 1280 matrix per sequence. The same routine extracts representations either from the pre-trained weights or from a fine-tuned backbone, so that frozen and fine-tuned representations are compared under an identical protocol.

In developability tasks, a single peptide sequence is encoded and reduced to 1280-dimensional descriptor by mean pooling across residue positions, which is cached and passed to a predictor head. In the binding affinity task, we followed Alsamkary et al. [2025] and encoded peptide and target as separate sequences. Reducing the two matrices and combining them is part of the affinity predictor rather than of extraction (Section 5.4.2); because that reduction is trained, it is the per-residue matrices that are cached for the affinity models rather than a pooled descriptor. 52 affinity measurements were excluded because their target sequences exceeded the model’s 1,022-token limit.

#### 5.3.3 Structure prediction models

##### MSA

MSAs were generated with the ColabFold MMseqs2 server [Mirdita et al., 2022] via the client vendored in Boltz-2, using iterative profile search [Steinegger and Söding, 2017]. Chains shorter than 50 residues were not searched, as short peptides return few or no detectable homologues.

##### Boltz-2

Boltz-2 (v2.2.1) [Passaro et al., 2025] was used to extract peptide structure representations. Its trunk embeds the tokenised input and refines it through an MSA module and a pairformer stack, producing a single (per-token) representation 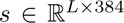 and a pairwise (per-token-pair) representation 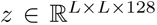, which the downstream diffusion and confidence modules consume. Peptide structures were predicted in single-sequence (MSA-free) mode with each peptide encoded as one protein chain, using three recycling iterations, 200 denoising steps and one diffusion sample per peptide.

For property prediction we discarded the diffusion and confidence modules and treated the trunk as a structure-aware sequence encoder, with five recycling iterations; for the kinetic stability dataset a recycling sweep showed no AUROC difference beyond run-to-run noise between zero and five iterations, so zero iterations were used to reduce cost by approximately fivefold per epoch. A prediction head then masked-mean-pooled the pairwise representation over valid token pairs to a single 128-dimensional vector and passed it through a three-layer MLP (128 → 128 → 64 → 1) with ReLU activations, emitting a logit for classification tasks or a scalar for regression tasks.

#### 5.3.4 Inverse folding

ProteinMPNN was used to obtain structure-aware inverse folding representations and sequence-structure compatibility scores [Dauparas et al., 2022]. Boltz-2 predicted structures were parsed to extract per-residue backbone coordinates. A proximity graph was built over the min(48, L) nearest neighbours of each residue by *C*α - *C*α distance, so for peptides shorter than 48 residues, the graph is effectively fully connected. Edges were featurised as radial basis expansions of all inter-atomic backbone distances plus relative sequence position. Per-residue embeddings were taken as the node features after the three message-passing encoder layers. For the perplexity analysis, sequence-structure compatibility was scored with the full encoder-decoder pass: the peptide sequence was integer-encoded over the 20 canonical amino acids plus an unknown token, embedded, and supplied to the autoregressive decoder, and the score was taken as the negative log-likelihood of that sequence averaged over residues. The score was averaged over 100 independently sampled random decoding orders and is reported as perplexity, the exponential of that average. The pre-trained ProteinMPNN model was used without fine-tuning (vanilla v_48_020 checkpoint, 48 edges, trained with 0.2 Å Gaussian backbone noise), with no backbone noise added at inference (σ = 0).

#### 5.3.5 Multimodal fusion of ESM2 and Boltz-2

For a peptide x of length N, the two models produce embeddings

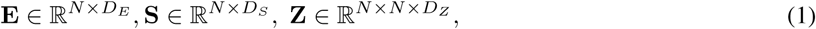

with D*_E_* = 1280, *D_S_* = 384 and *D_Z_* = 128. Boltz-2 is frozen in every fused model; ESM2 is either frozen or fully fine-tuned. For all three fusion strategies, the pair representation is reduced to a per-residue interaction profile by averaging over its second axis, and each modality is linearly projected to a common width *D* = 128,

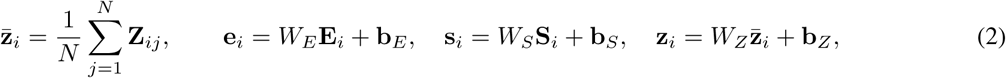

for i = 1, . .., *N*, with 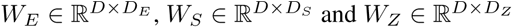 .

The strategies differ only in how the three projections are combined into a fused per-residue feature 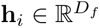,

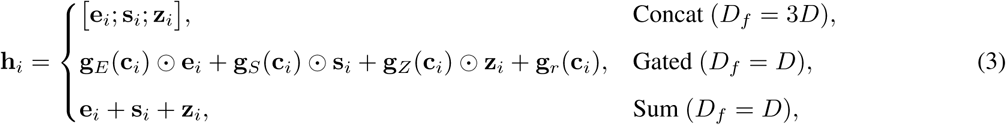

where 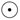 is the element-wise product and, for the gated variant Gu et al. [2026], the context vector and gates are

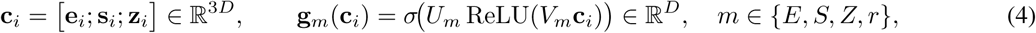

with bias-free 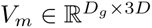 and 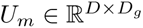, and σ the logistic sigmoid. The four gate networks share this form but not their weights, and **g***_r_* is an ungated residual term. Finally, the fused features are mean-pooled over the residue axis using the padding mask 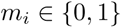 and passed to an MLP head.

### 5.4 Predictive Architectures and Training

#### 5.4.1 Developability Prediction

##### Classical machine learning baselines

For analysis of the peptide chemical space, descriptors that were non-constant and observed for at least half of the rows were retained, and the table was standardised to zero mean and unit variance before projection onto two partial least squares latent variables (PLS). Separability was scored out of fold by 5-fold cross-validation (stratified for the classification tasks), with standardisation and the PLS fit inside each fold. The classical predictive baseline was a gradient-boosted tree ensemble (XGBoost) on the same descriptors. Hyperparameters were selected by 5-fold grid search on the training split over the number of estimators (50, 100), the learning rate (0.05, 0.1, 0.2) and the maximum tree depth (3, 5), scoring accuracy for classification and negative mean absolute error for regression; all other settings were library defaults. The best configuration was refit on the full training split and evaluated once on the held-out test split.

##### ESM2

ESM2 was fine-tuned end to end for peptide property prediction. Every parameter of the encoder was updated together with those of the prediction head, without adapters or frozen layers. Sequences were tokenised with the ESM2 tokeniser and padded or truncated to a per-dataset maximum length chosen to cover that dataset’s length distribution. A fixed-length sequence representation was obtained by mean pooling the final hidden layer over non-padding tokens, followed by dropout (rate 0.1) and a linear head emitting two logits for classification tasks or a single scalar for regression tasks.

Classification used cross-entropy loss weighted by inverse class frequency, computed on the training split, and regression used mean squared error. Optimisation used AdamW (β_1_ = 0.9, β_2_ = 0.999, *∈* = 1 x 10*^-^*^8^) with a peak learning rate of 2 x 10*^-^*^5^, weight decay 0.01 and a batch size of 128. The learning rate was warmed up linearly over the first 10% of optimiser steps and then annealed to 1 x 10*^-^*^7^ by a cosine schedule, updated at every step. Training ran for at most 10 epochs with early stopping (patience 3 epochs) on validation AUROC for classification and on validation Spearman coefficient for regression, and the checkpoint achieving the best value of the monitored metric was retained for evaluation.

##### Boltz-2

Boltz-2 is too large to fine-tune fully on datasets of this size, and its pairwise track scales quadratically in sequence length, so we adapted it with Low-Rank Adaptation (LoRA) [Hu et al., 2022]. Adapters were injected into the last two pairformer layers, targeting all attention and triangular-update projections within each layer, 16 linear projections per layer in total. Each targeted projection with frozen weight *W* was replaced by 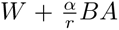, with 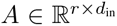 initialised from a Kaiming uniform distribution and 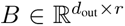 initialised to zero, so that the adapted model is numerically identical to the pre-trained one at initialisation. We used rank *r* = 8, scaling factor α = 16 and no LoRA dropout. All remaining trunk parameters were frozen and the trunk was kept in evaluation mode, so gradients flowed only to the adapter matrices and the prediction head; after training the adapters can be merged into the base weights, leaving inference cost unchanged.

Optimisation used AdamW with weight decay 1 x 10*^-^*^5^ and two parameter groups, the prediction head at a learning rate of 1 x 10*^-^*^4^ and the LoRA adapters at the lower rate of 5 x 10*^-^*^5^, with gradients clipped to a global norm of 1.0 and bfloat16 mixed precision. The learning rate was reduced by a factor of 0.5 after five epochs without improvement in the monitored validation metric. Focal loss was used (γ = 2, α = 0.25) with label smoothing 0.1 for the class-imbalanced classification datasets, binary cross-entropy for the class-balanced permeability dataset, and mean squared error for the continuous thermodynamic stability label. Batch size was set per dataset from the peptide length distribution, because the pairwise representation grows quadratically with sequence length: 32 for the aggregation and kinetic stability datasets, 12 for the permeability and thermodynamic stability datasets, and 8 for the non-fouling dataset. Training ran for at most 200 epochs with early stopping on validation AUROC (patience 10 epochs, minimum improvement 0.001) for classification and on validation loss (patience 15 epochs) for regression, retaining the best checkpoint by the monitored metric.

#### 5.4.2 Binding Affinity Prediction

##### ESM2

Each chain’s per-residue matrix (Section 5.3) is reduced to a single 1280-dimensional vector by attention pooling ([Alsamkary et al., 2025], Algorithm 1): a linear projection scores every residue, the scores are normalised over the sequence with padding excluded, and the residue representations are averaged under those weights. One projection serves both chains. The two vectors are concatenated into a 2560-dimensional descriptor and passed to an MLP of two hidden layers of width 512 with GELU activations and dropout, followed by a linear projection to the predicted affinity. The trained parameters are therefore only the pooling projection and the MLP.

###### Algorithm 1

Attention pooling

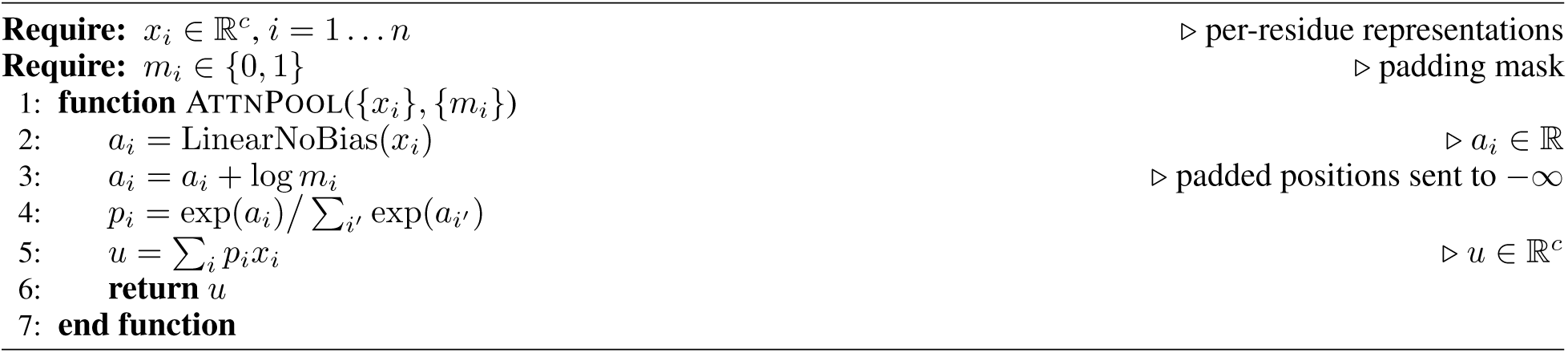

##### Boltz-2

Three of the affinity models are built on Boltz-2 [Passaro et al., 2025] and differ only in what sits between the pair representation of the cropped complex and the predicted affinity. in BOLTZ2 the pair representation is used as input to an MLP (called AffinityHeadsTransformer in the original code); the trunk is frozen and its output cached, so the head is the only trained component. AM (LIGAND) is the full affinity module with its released weights and no re-training. AM (PEPTIDE) is the same architecture trained on our dataset from random initialisation.

Inputs are constructed as in Passaro et al. [2025]: the complex is folded once, its predicted coordinates are used to crop it to the 256 tokens nearest the peptide (at most 200 of them from the target), and the affinity pass is run on that crop with five recycling steps. Anchor ordering, neighbourhood expansion and termination conditions in the cropper are unchanged, and all budgets retain their native values (Table 3). Two changes are required, both following from submitting the binder as a protein chain rather than as a SMILES ligand. The peptide is tokenised per residue rather than per atom, and binder tokens are identified by the affinity chain mask rather than by mol_type == NONPOLYMER, which is empty for a protein chain and would otherwise apply the receptor token cap to the binder as well as to the target. The second substitution propagates to the interface masks used downstream.

The module itself is otherwise unmodified (Algorithm 2). It conditions the pair representation on the token embeddings of the pretrained input embedder and on a distogram of the predicted geometry, refines it through four pairformer layers and passes it to the head; we refer to Passaro et al. [2025] for the definition of each stage. The geometry is taken from the highest-ipTM structure among the five sampled for the crop. Boltz-2 averages two trained variants of the module, one fit on more data for more epochs with a deeper pairformer; given the size of our dataset we use only the smaller variant.

###### Algorithm 2

Modified Boltz-2 affinity module

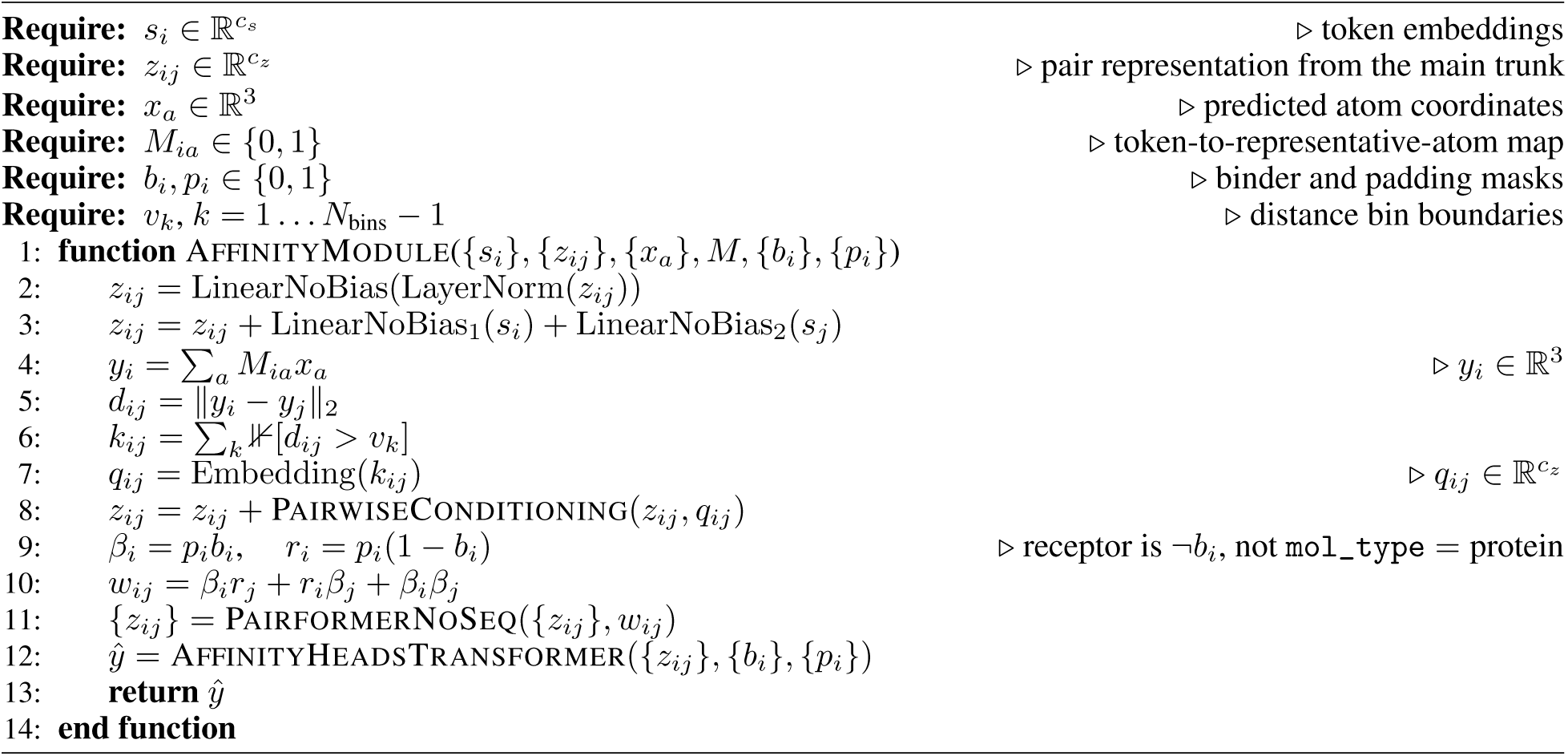

##### Batch construction

Following Passaro et al. [2025], our training objective supervises the relative differences between predictions rather than absolute values alone. We adopted this approach to control for the technical noise introduced by differing assay formats and laboratory protocols across publications. Consequently, this difference-based supervision dictated our batching strategy, requiring all samples within a batch to be drawn from the same experiment to ensure direct comparability. Measurements are partitioned into pools sharing PubMed identifier and measurement type. A publication may report against several target proteins, and about half of the training pools span more than one target sequence. Differences within a pool are therefore not always differences between peptides competing for the same protein. Measurements carrying no PubMed identifier were removed. Pools left with a single measurement were also discarded. During training, at each epoch pools are shuffled and sliced into batches of at most five, and the resulting batches are shuffled before being presented. Reshuffling within a pool means a different member falls into the short final batch on each pass, so no measurement is systematically under-represented, and every retained measurement is seen exactly once per epoch. Maximal batch size (5) is bounded by available memory and by the number of measurements sharing a publication.

##### Loss

Predictions are supervised by two terms, both using the Huber penalty *H_δ_*, which is quadratic for residuals below *δ* and linear beyond it. The same *δ* is used for both, although a residual of the second is a difference of differences. Labels are expressed as log_10_(µM), so a unit residual corresponds to one order of magnitude in concentration. The first term penalises the error on each predicted affinity,

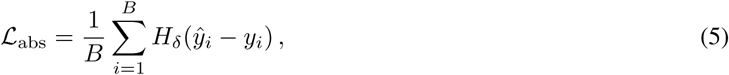

and the second penalises the error on every relative affinity within a batch of B ≥ 2 measurements,

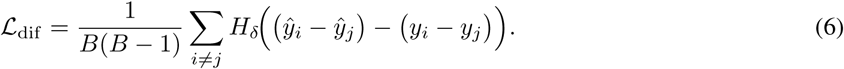

The total loss is

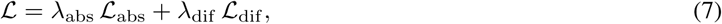

with λ_abs_ = 0.2, λ_dif_ = 1.8 and *δ* = 0.5, so the pairwise term carries nine times the weight of the absolute one.

##### Optimisation

Both representations are precomputed with frozen encoders and cached, so that training operates on a fixed set of features: the ESM2 encoder is not updated, and pair representations are extracted once from a frozen Boltz-2 trunk. What is trained on top of them differs by model: the pooling projection and MLP for ESM2, the affinity head alone for the pair representation, and the whole affinity module when that is used in place of the head. The module is trained from random initialisation; we also report it with Boltz-2’s released affinity weights and no training at all.

Models were trained with AdamW for a maximum of 30 epochs, with a warmup–cosine learning-rate schedule and early stopping on the validation pairwise-difference loss (patience 10); the checkpoint minimising that quantity was retained. Hyperparameters are given in Table 3.

##### Evaluation

For each panel we report Spearman’s *ρ*, Kendall’s τ*_b_*, Pearson’s r, and mean absolute error in log_10_units, together with the number of measurements and the mean pairwise sequence similarity of the peptides.

#### 5.4.3 Experimental setup and computational resources

To ensure strict reproducibility and standardize computational performance, all model training, fine-tuning, and structure prediction inference (including ESM2, Boltz-2, and ProteinMPNN) were executed on a single NVIDIA H100 80 GB SXM GPU. A global random seed (42) was fixed across all experimental pipelines, governing dataset partitioning, batch sampling, and model weight initialization. Where applicable, neural network optimization utilized bfloat16 mixed precision to maximize memory efficiency without compromising numerical stability.

## 6 Data and Code Availability

Datasets used in this study, model weights and inference code are available at https://huggingface.co/datasets/InstaDeepAI/PeptideScorer and https://huggingface.co/InstaDeepAI/PeptideScorer.

**Table 3:**
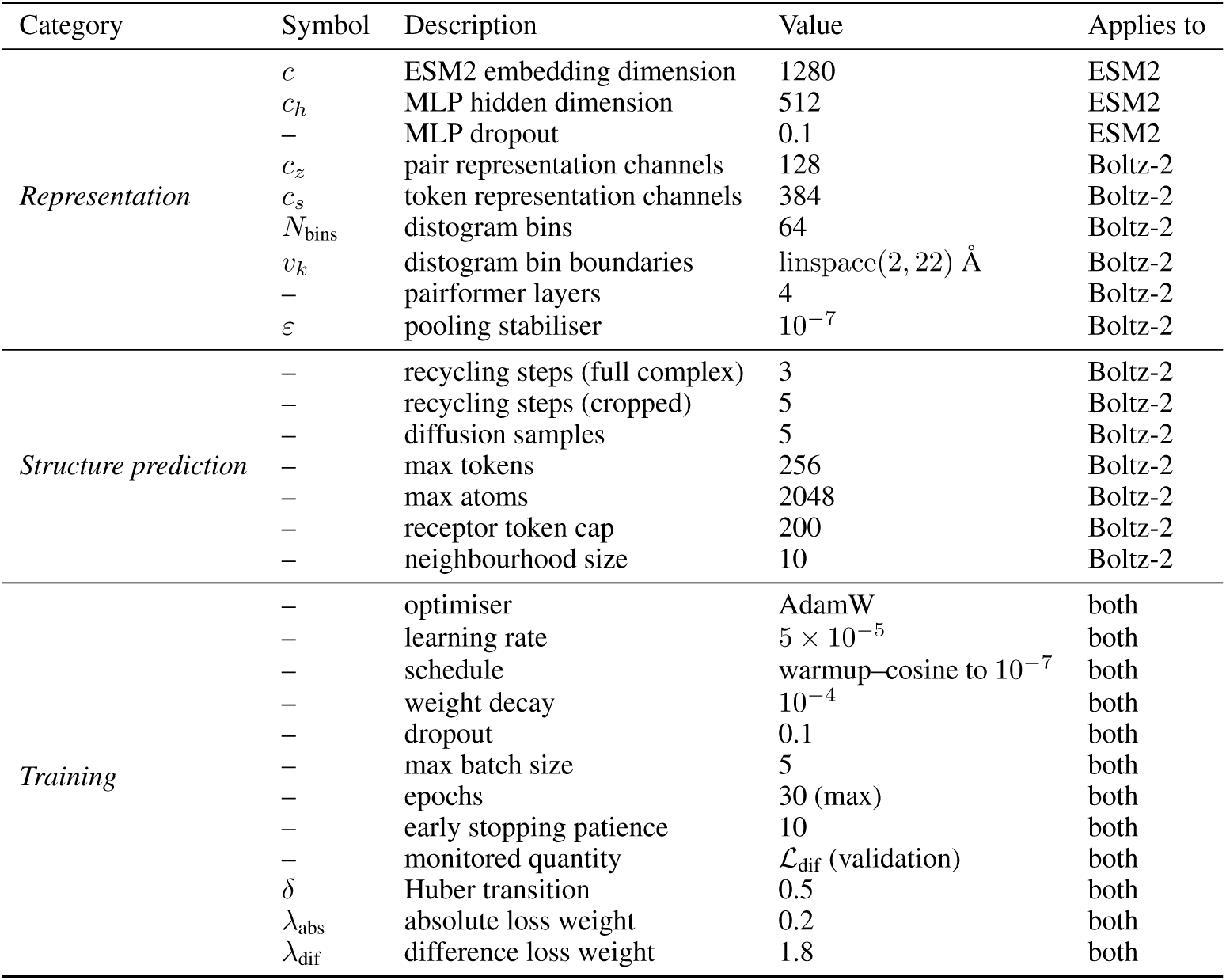
Hyperparameters for binding affinity prediction.

## Author Contributions

Q.T.: Methodology, Software, Validation, Formal Analysis, Data Curation, Writing - Original Draft, Writing - Review & Editing, Visualization. A.A.: Methodology, Software, Data Curation, Investigation. A.B.: Conceptualization, Writing - Original Draft, Supervision. Y.F: Conceptualization, Supervision. J.P.E.: Software (Lead), Writing - Review & Editing. C.Z.: Writing - Review & Editing, Supervision. F.S.: Conceptualization, Methodology, Software, Validation, Formal Analysis, Data Curation, Writing - Original Draft, Writing - Review & Editing, Visualization, Supervision.

## Footnotes

1 https://github.com/PistoiaHELM/HELMMonomerSets

2 https://www.rdkit.org

## References

Markus Muttenthaler, Glenn F. King, David J. Adams, and Paul F. Alewood. Trends in peptide drug discovery. Nature Reviews Drug Discovery, 20(4):309–325, 2021. doi: 10.1038/s41573-020-00135-8.

Martin Pacesa, Lennart Nickel, Christian Schellhaas, Joseph Schmidt, Ekaterina Pyatova, Lucas Kissling, Patrick Barendse, Jagrity Choudhury, Srajan Kapoor, Ana Alcaraz-Serna, Yehlin Cho, Kourosh H. Ghamary, Laura Vinué, Brahm J. Yachnin, Andrew M. Wollacott, Stephen Buckley, Adrie H. Westphal, Simon Lindhoud, Sandrine Georgeon, Casper A. Goverde, Georgios N. Hatzopoulos, Pierre Gönczy, Yannick D. Muller, Gerald Schwank, Daan C. Swarts, Alex J. Vecchio, Bernard L. Schneider, Sergey Ovchinnikov, and Bruno E. Correia. One-shot design of functional protein binders with bindcraft. Nature, 646:483–492, 10 2025. ISSN 14764687. doi: 10.1038/s41586-025-09429-6.

Hannes Stark, Felix Faltings, MinGyu Choi, Yuxin Xie, Eunsu Hur, Timothy O’Donnell, Anton Bushuiev, Talip Uçar, Saro Passaro, Weian Mao, Mateo Reveiz, Roman Bushuiev, Tally Portnoi, Tomáš Pluskal, Josef Sivic, Karsten Kreis, Arash Vahdat, Shamayeeta Ray, Jonathan T. Goldstein, Andrew Savinov, Jacob A. Hambalek, Anshika Gupta, Diego A. Taquiri-Diaz, Yaotian Zhang, Samuel J. Snyder, A. Katherine Hatstat, Angelika Arada, Nam Hyeong Kim, Haoyu Fan, Ethel Tackie-Yarboi, Dylan Boselli, Lee Schnaider, Chang C. Liu, Gene-Wei Li, Denes Hnisz, David M. Sabatini, William F. DeGrado, Jeremy Wohlwend, Gabriele Corso, Regina Barzilay, and Tommi Jaakkola. Boltzgen: Toward universal binder design. *bioRXiv*, 11 2025. doi: 10.1101/2025.11.20.689494. URL http://biorxiv.org/lookup/doi/10.1101/2025.11.20.689494.

Stephen A. Rettie, David Juergens, Victor Adebomi, Yensi Flores Bueso, Qinqin Zhao, Alexandria N. Leveille, Andi Liu, Asim K. Bera, Joana A. Wilms, Alina Üffing, Alex Kang, Evans Brackenbrough, Mila Lamb, Stacey R. Gerben, Analisa Murray, Paul M. Levine, Maika Schneider, Vibha Vasireddy, Sergey Ovchinnikov, Oliver H. Weiergräber, Dieter Willbold, Joshua A. Kritzer, Joseph D. Mougous, David Baker, Frank DiMaio, and Gaurav Bhardwaj. Accurate de novo design of high-affinity protein-binding macrocycles using deep learning. Nature Chemical Biology, 6 2025. ISSN 1552-4450. doi: 10.1038/s41589-025-01929-w. URL https://www.nature.com/articles/s41589-025-01929-w.

Latent Labs Team, Alex Bridgland, Jonathan Crabbé, Henry Kenlay, Daniella Pretorius, Sebastian M. Schmon, Agrin Hilmkil, Rebecca Bartke-Croughan, Robin Rombach, Michael Flashman, Tomas Matteson, Simon Mathis, Alexander W. R. Nelson, David Yuan, Annette Obika, and Simon A. A. Kohl. Latent-x: An atom-level frontier model for de novo protein binder design. *arXiv*, 7 2025. URL http://arxiv.org/abs/2507.19375.

Giacomo Rossino, Emanuela Marchese, Giovanni Galli, Francesca Verde, Matteo Finizio, Massimo Serra, Pasquale Linciano, and Simona Collina. Peptides as therapeutic agents: Challenges and opportunities in the green transition era. Molecules, 28(20):7165, 2023. doi: 10.3390/molecules28207165.

Jianan Li, Keisuke Yanagisawa, Masatake Sugita, Takuya Fujie, Masahito Ohue, and Yutaka Akiyama. Cycpeptmpdb: a comprehensive database of membrane permeability of cyclic peptides. Journal of chemical information and modeling, 63(7):2240–2250, 2023.

Andrew L. Hopkins, Colin R. Groom, and Alexander Alex. Ligand efficiency: a useful metric for lead selection. Drug Discovery Today, 9:430–431, 5 2004. ISSN 13596446. doi: 10.1016/S1359-6446(04)03069-7. URL https://linkinghub.elsevier.com/retrieve/pii/S1359644604030697.

Jolene L Lau and Michael K Dunn. Therapeutic peptides: Historical perspectives, current development trends, and future directions. Bioorganic & medicinal chemistry, 26(10):2700–2707, 2018.

Vanessa Erckes, Massina Abderrahmane, Maud Jusot, Christian Steuer, and Rodrigo Ochoa. Peptide cheminformatics tools: making computational tasks accessible in peptide drug discovery. Drug Discovery Today, 31:104612, 3 2026. ISSN 13596446. doi: 10.1016/j.drudis.2026.104612. URL https://linkinghub.elsevier.com/retrieve/pii/S1359644626000176.

Zeming Lin, Halil Akin, Roshan Rao, Brian Hie, Zhongkai Zhu, Wenting Lu, Nikita Smetanin, Robert Verkuil, Ori Kabeli, Yaniv Shmueli, et al. Evolutionary-scale prediction of atomic-level protein structure with a language model. Science, 379(6637):1123–1130, 2023.

John Jumper, Richard Evans, Alexander Pritzel, Tim Green, Michael Figurnov, Olaf Ronneberger, Kathryn Tunyasuvunakool, Russ Bates, Augustin Žídek, Anna Potapenko, Alex Bridgland, Clemens Meyer, Simon A. A. Kohl, Andrew J. Ballard, Andrew Cowie, Bernardino Romera-Paredes, Stanislav Nikolov, Rishub Jain, Jonas Adler, Trevor Back, Stig Petersen, David Reiman, Ellen Clancy, Michal Zielinski, Martin Steinegger, Michalina Pacholska, Tamas Berghammer, Sebastian Bodenstein, David Silver, Oriol Vinyals, Andrew W. Senior, Koray Kavukcuoglu, Pushmeet Kohli, and Demis Hassabis. Highly accurate protein structure prediction with AlphaFold. Nature, 596(7873): 583–589, 2021. doi: 10.1038/s41586-021-03819-2.

Saro Passaro, Gabriele Corso, Jeremy Wohlwend, Mateo Reveiz, Stephan Thaler, Vignesh Ram Somnath, Noah Getz, Tally Portnoi, Julien Roy, Hannes Stark, David Kwabi-Addo, Dominique Beaini, Tommi Jaakkola, and Regina Barzilay. Boltz-2: Towards accurate and efficient binding affinity prediction. *bioRXiv*, 2025. doi: 10.1101/2025.06.14.659707.

Chakradhar Guntuboina, Adrita Das, Parisa Mollaei, Seongwon Kim, and Amir Barati Farimani. Peptidebert: A language model based on transformers for peptide property prediction. Journal of Physical Chemistry Letters, 14: 10427–10434, 11 2023. ISSN 19487185. doi: 10.1021/acs.jpclett.3c02398.

Ruochi Zhang, Haoran Wu, Chang Liu, Qian Yang, Yuting Xiu, Kewei Li, Ningning Chen, Yu Wang, Yan Wang, Xin Gao, and Fengfeng Zhou. Pepland: a large-scale pre-trained peptide representation model for a comprehensive landscape of both canonical and non-canonical amino acids. Briefings in Bioinformatics, 26, 7 2025. ISSN 1467-5463. doi: 10.1093/bib/bbaf367. URL https://academic.oup.com/bib/article/doi/10.1093/bib/bbaf367/8220755.

Xiaorong Tan, Qianhui Liu, Mengting Zhou, and Yanpeng Fang. pepadmet: A novel computational platform for systematic admet evaluation of peptides. Journal of Chemical Information and Modeling, 2026. doi: 10.1021/acs.jcim.5c02518.

Nikhil Shenoy, David Errington, Emmanuel Bengio, Kacper Kapusniak, Kerstin Klaeser, Yui Tik Pang, Vladimir Radenkovic, Prudencio Tossou, Therence Bois, Andrew Wedlake, and Francesco Di Giovanni. Nesso-1: Accelerating open-source binding affinity predictions. Technical report, Valence Labs, Recursion, July 2026. URL https://www.valencelabs.com/wp-content/uploads/2026/07/nesso1.pdf.

SandboxAQ. AQAffinity: Open-source structure-to-affinity built on OpenFold3. https://www.sandboxaq.com/post/aqaffinity-open-source-structure-to-affinity-built-on-openfold3-by-sandboxaq, 2026. Accessed: 2026-09-04.

Dan Ofer and Michal Linial. Profet: Feature engineering captures high-level protein functions. Bioinformatics, 31: 3429–3436, 12 2015. ISSN 14602059. doi: 10.1093/bioinformatics/btv345.

Yinuo Zhang, Sophia Tang, Tong Chen, Elizabeth Mahood, Sophia Vincoff, and Pranam Chatterjee. Peptiverse: A unified platform for therapeutic peptide property prediction. Nature Communications, 17:6819, 7 2026. ISSN 2041-1723. doi: 10.1038/s41467-026-74167-w. URL https://www.nature.com/articles/s41467-026-74167-w.

Josh Abramson, Jonas Adler, Jack Dunger, Richard Evans, Tim Green, Alexander Pritzel, Olaf Ronneberger, Lindsay Willmore, Andrew J. Ballard, Joshua Bambrick, Sebastian W. Bodenstein, David A. Evans, Chia Chun Hung, Michael O’Neill, David Reiman, Kathryn Tunyasuvunakool, Zachary Wu, Akvilé Žemgulyté, Eirini Arvaniti, Charles Beattie, Ottavia Bertolli, Alex Bridgland, Alexey Cherepanov, Miles Congreve, Alexander I. Cowen-Rivers, Andrew Cowie, Michael Figurnov, Fabian B. Fuchs, Hannah Gladman, Rishub Jain, Yousuf A. Khan, Caroline M.R. Low, Kuba Perlin, Anna Potapenko, Pascal Savy, Sukhdeep Singh, Adrian Stecula, Ashok Thillaisundaram, Catherine Tong, Sergei Yakneen, Ellen D. Zhong, Michal Zielinski, Augustin Žídek, Victor Bapst, Pushmeet Kohli, Max Jaderberg, Demis Hassabis, and John M. Jumper. Accurate structure prediction of biomolecular interactions with alphafold 3. Nature, 630:493–500, 6 2024. ISSN 14764687. doi: 10.1038/s41586-024-07487-w.

Qurat-ul-ain, Carlos Outeiral, Matteo Cagiada, Yee Whye Teh, and Charlotte M. Deane. Just add structure: Protein language models combined with structural equivariance excel at protein tasks. *bioRXiv*, 5 2026. doi: 10.64898/2026.05.28.728196. URL http://biorxiv.org/lookup/doi/10.64898/2026.05.28.728196.

Ziang Li and Yunan Luo. Generalizable and scalable protein stability prediction with rewired protein generative models. Nature Communications, 2025.

Lei Wang, Nanxi Wang, Wenping Zhang, Xurui Cheng, Zhibin Yan, Gang Shao, Xi Wang, Rui Wang, and Caiyun Fu. Therapeutic peptides: current applications and future directions. Signal Transduction and Targeted Therapy, 7(1):47, 2022. doi: 10.1038/s41392-022-00904-4.

Björn Mattsson and W. Patrick Walters. Identifying and addressing systematic data leakage in protein-ligand affinity benchmarks. *bioRxiv*, 6 2026. doi: 10.64898/2026.06.29.735309. URL http://biorxiv.org/lookup/doi/10.64898/2026.06.29.735309.

Lingle Wang, Yujie Wu, Yuqing Deng, Byungchan Kim, Levi Pierce, Goran Krilov, Dmitry Lupyan, Shaughnessy Robinson, Markus K. Dahlgren, Jeremy Greenwood, Donna L. Romero, Craig Masse, Jennifer L. Knight, Thomas Steinbrecher, Thijs Beuming, Wolfgang Damm, Edward Harder, Woody Sherman, Mark Brewer, Ron Wester, Mark Murcko, Leah Frye, Ramy Farid, Teng Lin, David L. Mobley, William L. Jorgensen, Bruce J. Berne, Richard A. Friesner, and Robert Abel. Accurate and reliable prediction of relative ligand binding potency in prospective drug discovery by way of a modern free-energy calculation protocol and force field. Journal of the American Chemical Society, 137(7):2695–2703, 2015. doi: 10.1021/ja512751q.

Mike Thompson, Mariano Martín, Trinidad Sanmartín Olmo, Chandana Rajesh, Peter K. Koo, Benedetta Bolognesi, and Ben Lehner. Massive experimental quantification allows interpretable deep learning of protein aggregation. Science Advances, 11:5111, 5 2025. ISSN 2375-2548. doi: 10.1126/sciadv.adt5111. URL https://www.science.org/doi/10.1126/sciadv.adt5111.

Cristina Chiva, Zahra Elhamraoui, Amanda Solé, Marc Serret, Mathias Wilhelm, and Eduard Sabidó. Assessment and prediction of human proteotypic peptide stability for proteomics quantification. Analytical Chemistry, 95(37): 13746–13749, 2023.

Jin Su, Chenchen Han, Yuyang Zhou, Junjie Shan, Xibin Zhou, and Fajie Yuan. Saprot: Protein language modeling with structure-aware vocabulary. *bioRxiv*, 2023.doi: 10.1101/2023.10.01.560349.

Ning Zhu, Yanyu Ming, Chengyun Zhang, Cao Sen, Chongyang Li, Jingjing Guo, and Hongliang Duan. Ppikb: A comprehensive knowledge base and analysis platform for protein–peptide interactions based on literature and patents. bioRxiv, pages 2025–06, 2025.

Barbara Zdrazil, Eloy Felix, Fiona Hunter, Emma J Manners, James Blackshaw, Sybilla Corbett, Marleen de Veij, Harris Ioannidis, David Mendez Lopez, Juan F Mosquera, Maria Paula Magarinos, Nicolas Bosc, Ricardo Arcila, Tevfik Kizilören, Anna Gaulton, A Patrícia Bento, Melissa F Adasme, Peter Monecke, Gregory A Landrum, and Andrew R Leach. The chembl database in 2023: a drug discovery platform spanning multiple bioactivity data types and time periods. Nucleic Acids Research, 52:D1180–D1192, 1 2024. ISSN 0305-1048. doi: 10.1093/nar/gkad1004. URL https://academic.oup.com/nar/article/52/D1/D1180/7337608.

Huaqing Liu, Peiyi Chen, Xiaochen Zhai, Ku-Geng Huo, Shuxian Zhou, Lanqing Han, and Guoxin Fan. Ppb-affinity: Protein-protein binding affinity dataset for ai-based protein drug discovery. Scientific data, 11(1):1316, 2024.

Ann K Nowinski, Fang Sun, Andrew D White, Andrew J Keefe, and Shaoyi Jiang. Sequence, structure, and function of peptide self-assembled monolayers. Journal of the American Chemical Society, 134:6000–6005, 3 2012. ISSN 0002-7863. doi: 10.1021/ja3006868. URL 10.1021/ja3006868.

Tianqi Chen and Carlos Guestrin. XGBoost: A scalable tree boosting system. In Proceedings of the 22nd ACM SIGKDD International Conference on Knowledge Discovery and Data Mining, pages 785–794. ACM, 2016. doi: 10.1145/2939672.2939785.

Yannan Bin, Daijun Zhang, Zhiyang Hu, Chungui Xu, and Yansen Su. Pepxml: Esm2-based extreme multilabel classification of pathogen-targeted antimicrobial peptides. Briefings in Bioinformatics, 26, 9 2025. ISSN 14774054. doi: 10.1093/bib/bbaf548.

Dea Gogishvili, Emmanuel Minois-Genin, Jan Van Eck, and Sanne Abeln. Patchprot: hydrophobic patch prediction using protein foundation models. Bioinformatics Advances, 4, 2024. ISSN 26350041. doi: 10.1093/bioadv/vbae154.

Sophie E. Jackson. How do small single-domain proteins fold? Folding and Design, 3:R81–R91, 8 1998. ISSN 13590278. doi: 10.1016/S1359-0278(98)00033-9. URL https://linkinghub.elsevier.com/retrieve/pii/S1359027898000339.

Justas Dauparas, Ivan Anishchenko, Nathaniel Bennett, Hua Bai, Robert J Ragotte, Lukas F Milles, Basile IM Wicky, Alexis Courbet, Rob J de Haas, Neville Bethel, et al. Robust deep learning–based protein sequence design using proteinmpnn. Science, 378(6615):49–56, 2022.

James King, Lewis Cornwall, Andrei Cristian Nica, James Day, Aaron Sim, Neil Dalchau, Lilly Wollman, and Joshua Meyers. On fine-tuning boltz-2 for protein-protein affinity prediction. *aRXiv*, 12 2025. URL http://arxiv.org/abs/2512.06592.

Hazem Alsamkary, Mohamed Elshaffei, Mohamed Soudy, Sara Ossman, Abdallah Amr, Nehal Adel Abdelsalam, Mohamed Elkerdawy, and Ahmed Elnaggar. Beyond simple concatenation: fairly assessing plm architectures for multi-chain protein-protein interactions prediction. arXiv preprint arXiv:2505.20036, 2025.

I. D. Kuntz, K. Chen, K. A. Sharp, and P. A. Kollman. The maximal affinity of ligands. Proceedings of the National Academy of Sciences, 96:9997–10002, 8 1999. ISSN 0027-8424. doi: 10.1073/pnas.96.18.9997. URL https://pnas.org/doi/full/10.1073/pnas.96.18.9997.

Tong Zhu, Shuai Cao, Pin-Chih Su, Ram Patel, Darshan Shah, Heta B. Chokshi, Richard Szukala, Michael E. Johnson, and Kirk E. Hevener. Hit identification and optimization in virtual screening: Practical recommendations based on a critical literature analysis. Journal of Medicinal Chemistry, 56(17):6560–6572, 2013. doi: 10.1021/jm301916b.

Jianyuan Deng, Zhibo Yang, Hehe Wang, Iwao Ojima, Dimitris Samaras, and Fusheng Wang. A systematic study of key elements underlying molecular property prediction. Nature Communications, 14, 12 2023. ISSN 20411723. doi: 10.1038/s41467-023-41948-6.

Matthew R. Masters, Amr H. Mahmoud, and Markus A. Lill. Investigating whether deep learning models for co-folding learn the physics of protein-ligand interactions. Nature Communications, 16(1):8854, 2025. doi: 10.1038/s41467-025-63947-5.

Ryan Park, Darren J. Hsu, C. Brian Roland, Maria Korshunova, Chen Tessler, Shie Mannor, Olivia Viessmann, and Bruno Trentini. Improving inverse folding for peptide design with diversity-regularized direct preference optimization. *bioRxiv*, 10 2024. URL http://arxiv.org/abs/2410.19471.

Gabriel J. Rocklin, Tamuka M. Chidyausiku, Inna Gober, Ash Ford, Alexander Groom, Patrick J. Hember, Hao Dou, Yoav Peleg, Brian P. Mendes, Fabian Schwetlick, Daniel J. Thompson, Francis Barry, Richard Bonneau, and David Baker. Global analysis of protein folding using massively parallel design, synthesis, and testing. Science, 357(6347): 168–175, 2017. doi: 10.1126/science.aan0693.

Tianhong Zhang, Hongli Li, Hualin Xi, Robert V. Stanton, and Sergio H. Rotstein. Helm: A hierarchical notation language for complex biomolecule structure representation. Journal of Chemical Information and Modeling, 52: 2796–2806, 10 2012. ISSN 1549960X. doi: 10.1021/ci3001925.

Gregory A. Landrum and Sereina Riniker. Combining ic50 or ki values from different sources is a source of significant noise. Journal of Chemical Information and Modeling, 64:1560–1567, 3 2024. ISSN 1549960X. doi: 10.1021/acs.jcim.4c00049.

Shafayat Ahmed, Muhit Islam Emon, Nazifa Ahmed Moumi, and Liqing Zhang. Fast-part: Fast and accurate data partitioning for biological sequence analysis. bioRxiv, pages 2024–11, 2024.

Martin Steinegger and Johannes Söding. Mmseqs2 enables sensitive protein sequence searching for the analysis of massive data sets. Nature biotechnology, 35(11):1026–1028, 2017.

Benjamin Buchfink, Klaus Reuter, and Hajk Georg Drost. Sensitive protein alignments at tree-of-life scale using diamond. Nature Methods, 18:366–368, 4 2021. ISSN 15487105. doi: 10.1038/s41592-021-01101-x.

Peter J.A. Cock, Tiago Antao, Jeffrey T. Chang, Brad A. Chapman, Cymon J. Cox, Andrew Dalke, Iddo Friedberg, Thomas Hamelryck, Frank Kauff, Bartek Wilczynski, and Michiel J.L. De Hoon. Biopython: Freely available python tools for computational molecular biology and bioinformatics. Bioinformatics, 25:1422–1423, 6 2009. ISSN 13674803. doi: 10.1093/bioinformatics/btp163.

Alex S. Holehouse, Rahul K. Das, James N. Ahad, Mary O.G. Richardson, and Rohit V. Pappu. Cider: Resources to analyze sequence-ensemble relationships of intrinsically disordered proteins. Biophysical Journal, 112:16–21, 1 2017. ISSN 15420086. doi: 10.1016/j.bpj.2016.11.3200.

Mauno Vihinen, Esa Torkkila, and Pentti Riikonen. Accuracy of protein flexibility predictions. Proteins: Structure, Function, and Bioinformatics, 19(2):141–149, 1994. doi: 10.1002/prot.340190207. URL https://onlinelibrary.wiley.com/doi/abs/10.1002/prot.340190207.

Liu Yang, Suqi Cao, Lei Liu, Ruixin Zhu, and Dingfeng Wu. cyclicpeptide: a python package for cyclic peptide drug design. Briefings in Bioinformatics, 26, 1 2025. ISSN 14774054. doi: 10.1093/bib/bbae714.

Milot Mirdita, Konstantin Schütze, Yoshitaka Moriwaki, Lim Heo, Sergey Ovchinnikov, and Martin Steinegger. Colabfold: making protein folding accessible to all. Nature Methods, 19:679–682, 6 2022. ISSN 15487105. doi: 10.1038/s41592-022-01488-1.

Chunbin Gu, Zijun Gao, Mutian He, Jingjie Zhang, Haipeng Wen, Zihao Luo, Xiaorui Wang, Hanqun Cao, Jiajun Bu, Chang-Yu Hsieh, and Pheng Ann Heng. Bi-team: A unified cross-scale representation learning framework for chemically modified biomolecules. *arXiv*, 3 2026. URL http://arxiv.org/abs/2603.01873.

Edward J Hu, Yelong Shen, Phillip Wallis, Zeyuan Allen-Zhu, Yuanzhi Li, Shean Wang, Lu Wang, Weizhu Chen, et al. Lora: Low-rank adaptation of large language models. ICLR, 1(2):3, 2022.

