## Supplementary Information for "Chemical Descriptors and Deep Learning Embeddings for Scoring *de novo* Peptide Designs"

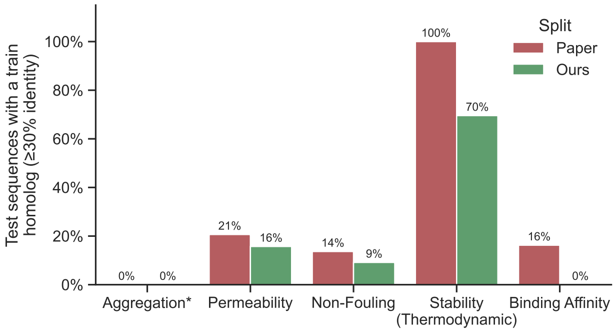


Figure S1: Train-test homology leakage across datasets under the published splits (Paper) versus our leakage-aware Fast-Part splits (Ours). Each bar shows the percentage of test sequences with at least one training-set homologue at ≥30% sequence identity. *Aggregation peptides ( 20 aa) are near the sensitivity floor of the homology audit, so its 0% leakage partly reflects limited detectability at this length, not only a clean split. The analysis is limited to dataset with data splits available in the literature.

| ESM2 | Boltz-2 | Fusion | Aggregation | | Permeability | | Non-Fouling | | Stability (Kinetic) | | Stability (Thermodynamic) | |
|---|---|---|---|---|---|---|---|---|---|---|---|---|
| | | | AUROC | AUPRC | AUROC | AUPRC | AUROC | AUPRC | AUROC | AUPRC | Spearman | Pearson |
| *Unimodal baselines* | | | | | | | | | | | | |
| Frozen | – | – | 68.9 | 40.7 | 92.8 | 94.8 | 84.0 | 67.9 | 69.9 | 88.7 | 0.577 | 0.579 |
| Full FT | – | – | **73.9** | **51.8** | 91.9 | 94.5 | 84.9 | 70.3 | **79.8** | **93.9** | 0.717 | 0.741 |
| – | Frozen | – | 69.6 | 42.2 | 92.1 | 94.6 | 83.9 | 68.8 | 75.7 | 92.4 | 0.603 | 0.611 |
| – | LoRA FT | – | 69.7 | 42.4 | 92.5 | 94.9 | 84.6 | 70.6 | 75.8 | 92.4 | 0.610 | 0.615 |
| *Fused* | | | | | | | | | | | | |
| Frozen | Frozen | Sum | 70.8 | 45.0 | 93.0 | 95.1 | 84.8 | 70.1 | 75.6 | 92.1 | 0.649 | 0.681 |
| Frozen | Frozen | Concat | 71.0 | 45.8 | 94.2 | 96.0 | **85.0** | **70.7** | 76.2 | 92.4 | 0.664 | 0.699 |
| Frozen | Frozen | Gated | 71.4 | 47.5 | 94.4 | 95.9 | 84.4 | 69.0 | 76.7 | 92.7 | 0.684 | 0.706 |
| *Fused (Fine-tuned ESM2)* | | | | | | | | | | | | |
| Full FT | Frozen | Sum | 73.2 | 49.4 | 93.7 | 95.6 | 84.2 | 67.7 | 78.5 | 93.0 | 0.754 | 0.770 |
| Full FT | Frozen | Concat | 72.9 | 48.4 | 94.2 | 96.2 | 84.8 | 69.8 | 79.3 | 93.7 | **0.770** | **0.782** |
| Full FT | Frozen | Gated | 72.1 | 45.7 | **94.8** | **96.7** | 83.9 | 67.4 | 79.0 | 93.4 | 0.749 | 0.755 |

Table S1: Ablation study of ESM2 and Boltz-2 encoder configurations and fusion strategies across the peptide developability datasets. AUROC and AUPRC (%) are reported as classification performance metrics. Bold indicates the best performing configuration per dataset; underline indicates the second best. Rows are grouped by ESM2 training regime: unimodal baselines, frozen-ESM2 fusion, and full fine-tuned ESM2 fusion. Fine-tuning of Boltz-2 was not tested on fused models due to compute limitations.

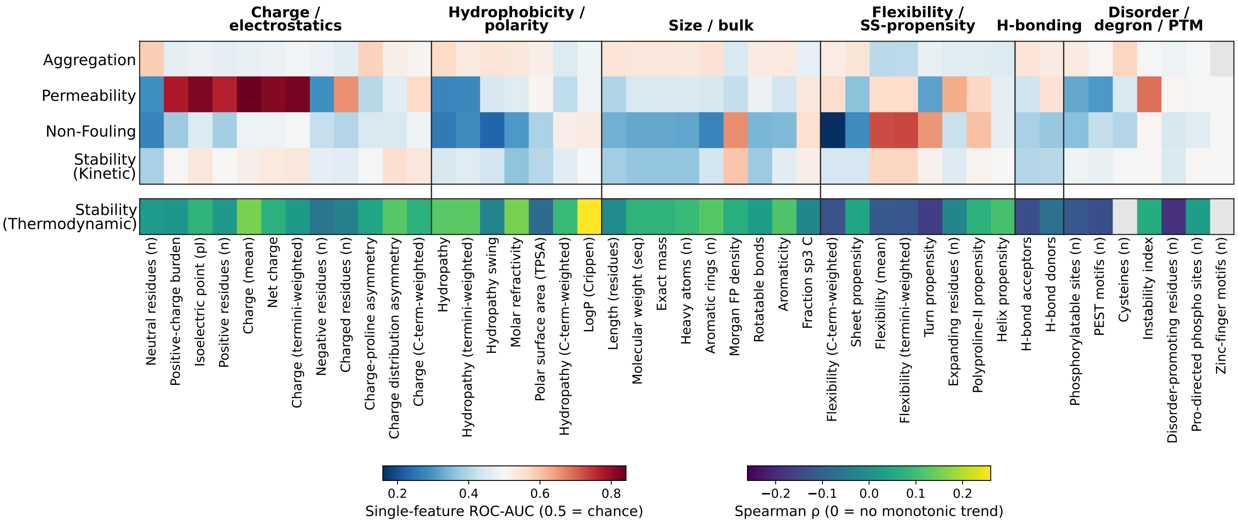


Figure S2: Individual predictive power of physicochemical descriptors across the five peptide tasks. Each cell reports how well a single descriptor, on its own, separates one task's label (curated data, peptides $\leq$100 residues). Descriptors (columns) are grouped into physicochemical families and ordered within each family by mean discriminative power across the four classification tasks. For the four binary tasks (top block) the cell value is the univariate ROC-AUC against the label, on a diverging blue-white-red scale centred at 0.5 (chance). Red (AUC $> 0.5$) marks descriptors higher in the positive class, blue (AUC $< 0.5$) higher in the negative class: positive/negative denote nucleating/non-nucleating (Aggregation), permeable/non-permeable (Permeability), non-fouling/fouling (Non-Fouling), and stable/unstable (Stability (Kinetic)). Stability (Thermodynamic), a continuous label, is shown as a separate strip (bottom) using Spearman's $\rho$ between descriptor and stability score: $\rho > 0$ (yellow) increases with, $\rho < 0$ (purple) decreases with, and $\rho \approx 0$ (green) is unrelated to predicted thermodynamic stability. Grey cells denote constant (undefined) descriptors.

| Model | Trainable params | Epochs | min/epoch | CPUs | RAM | GPU | Training time |
|---|---|---|---|---|---|---|---|
| Classical ML (XGBoost, 5-fold grid search) | — ($\sim$45 features) | — | — | 11 | 18 GB | None (CPU) | <1 min |
| ESM2-650M full fine-tune + head | 652 M (100%) | $\sim$6 | $\sim$0.5 | 16 | 64 GB | 1× H100 80 GB | $\sim$3 min |
| ESM2 frozen + Boltz-2 fusion | 361 K (0.06%) | $\sim$18 | $\sim$0.9 | 8 | 32 GB | 1× H100 80 GB | $\sim$17 min |
| ESM2 full + Boltz-2 fusion | 652 M (100%) | $\sim$18 | $\sim$1.4 | 16 | 64 GB | 1× H100 80 GB | $\sim$25 min |
| Boltz-2 LoRA (rank 8) + head | 127 K (102 K LoRA + 25 K head) | $\sim$40 | $\sim$57 | 16 | 64 GB | 1× H100 80 GB | $\sim$38 h |

Table S2: Approximate training time consumption per model on the non-fouling dataset. Hardware used: 2× Intel Xeon Platinum 8480+ (Sapphire Rapids, 224 threads, $\leq$3.8 GHz), $\sim$2 TB DDR5, 1× NVIDIA H100 80 GB HBM3 (700 W). Each job is cgroup-limited to the CPUs/RAM shown. Classical row measured locally: MacBook Pro 14″ (Apple M3 Pro, 11 cores 5P+6E $\sim$4.05 GHz, 18 GB LPDDR5).

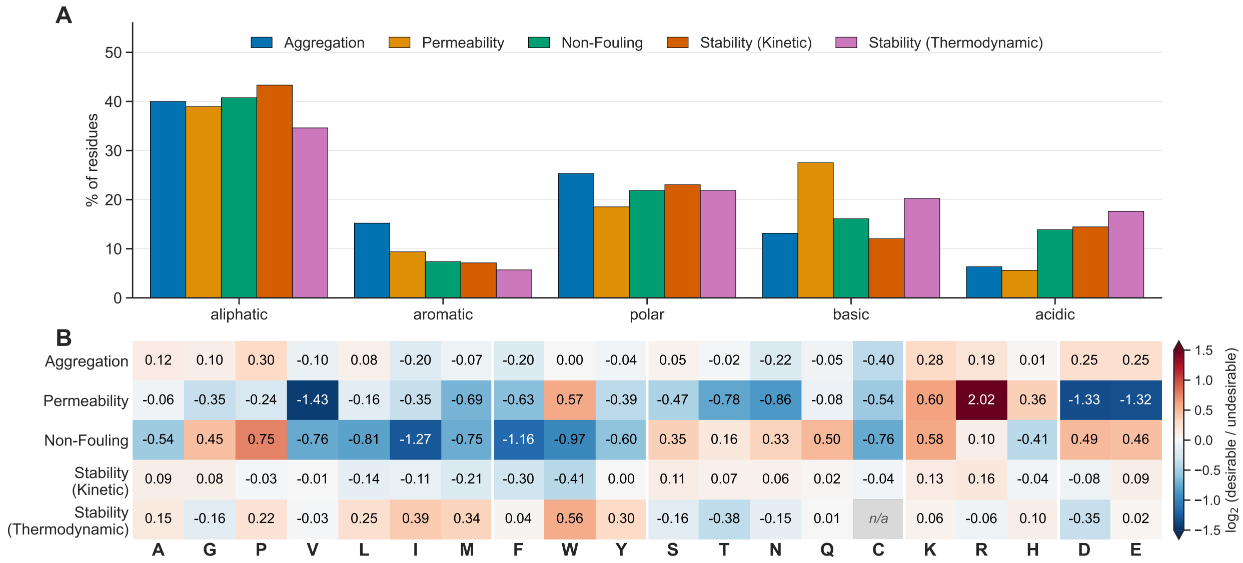


Figure S3: Amino-acid composition across the five developability datasets. (A) Residue composition of each dataset, as the percentage of residues falling in each of five side-chain families (aliphatic A, G, P, V, L, I, M; aromatic F, W, Y; polar S, T, N, Q, C; basic K, R, H; acidic D, E). Percentages are means of per-peptide residue frequencies and sum to 100% within each dataset. (B) Per-residue class enrichment, as $\log_2$ of the ratio of mean per-peptide residue frequency between the two label classes. Every row is oriented towards the developable outcome, named in parentheses beside the dataset, so positive (red) means the residue is enriched in peptides with the desirable property. For the one continuous task (thermodynamic stability) the classes are the top and bottom tertiles of the stability score. Grey *n/a* marks a residue absent from the dataset. Normalisation to residue counts per peptide is perfomed before comparing classes to make statistics insensitive to peptide length.

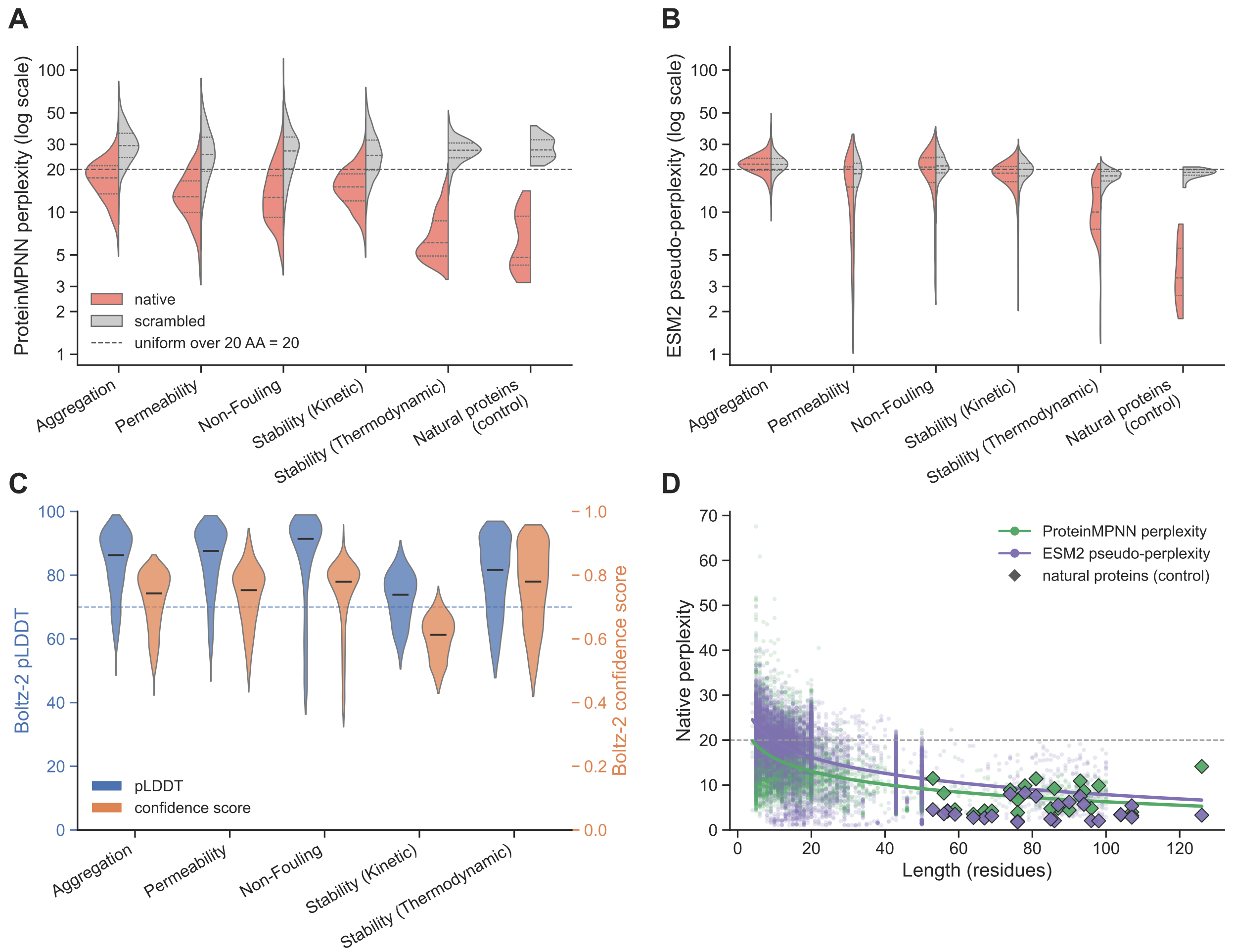


Figure S4: ESM2 and MPNN perplexity analysis and structure confidence metrics from Boltz-2. (A, B) ProteinMPNN perplexity and EMS2 pseudo-perplexity for each peptide's native sequence (red) versus a shuffled version of the same sequence threaded onto the same backbone (grey), with a positive control of 24 small natural proteins. The dashed line at 20 is the uniform-amino-acid baseline. (C) Per-peptide Boltz-2 confidence for 1500 folded peptides per dataset: pLDDT (blue, left axis) and the aggregate confidence score (orange, right axis). (D) Perplexity versus chain length, across the combined datasets (circles) and the natural protein controls (diamonds). Non-fouling peptides are capped at 100 aa. Control protein benchmark (from Jackson [1998]), 24 small single-domain proteins: 1LMB (P03034), 2ABD (P07107), 1HRC (P00004), 1YCC (P00044), 2AIT (P01092), 1CSP (P32081), 1MJC (P0A9X9), 1SHG (P07751), 1SRL (P00523), 1PKS (P27986), 1SHF (P06241), 1WIT (Q23551), 1TEN (P24821), 1TTF (P02751), 1COA (P01053), 1PBA (P09955), 1ARR (P03050), 1UBQ (P0CG48), 2PTL (Q53291), 1URN (P09012), 1HDN (P0AA04), 1FKB (P62942), 1APS (P00818), 2VIK (P02640).

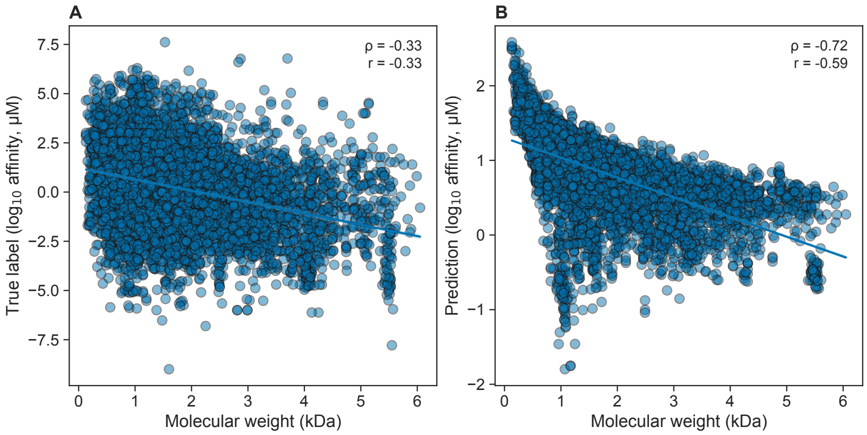


Figure S5: Confounding effect of MW on binding affinity predictions. (A) Linear regression of measured labels vs peptide molecular weights. (B) Linear regression of measured labels vs peptide molecular weights. Spearman $\rho$, Pearson's r and number of labels (n) are reported.

### References

Sophie E. Jackson. How do small single-domain proteins fold? *Folding and Design*, 3:R81–R91, 8 1998. ISSN 13590278. doi: 10.1016/S1359-0278(98)00033-9. URL https://linkinghub.elsevier.com/retrieve/pii/S1359027898000339.
